# Ancient MAX Effector Variants of a Fungal Pathogen Evade Apoplastic Immunity in Apple

**DOI:** 10.64898/2025.12.25.695362

**Authors:** Mélanie Sannier, Sirine Benmamar, Jérôme Collemare, Christophe Lemaire, Valérie Caffier, Clémentine Duplaix, Jason Shiller, Eglantine Mathieu-Bégné, Alice Vayssières, Henk J. Schouten, Carl H. Mesarich, Bruno Le Cam, Maël Baudin

## Abstract

*Venturia inaequalis* is an Ascomycota fungus responsible for apple scab, the main disease in apple orchards. During infection, the pathogen colonizes the subcuticular space and secretes effectors to promote virulence. To mitigate disease impact, resistance genes such as *Rvi6*, which encodes a membrane-localized receptor-like protein, have been introduced into apple cultivars. However, over the past three decades, *Rvi6*-mediated resistance has been circumvented in orchards, with the emergence of virulent *V. inaequalis* strains. Through comparative genomics analyses and functional validation by complementation, we identified *AvrRvi6* as the fungal determinant that activates *Rvi6*-mediated immunity. Screening 122 *V. inaequalis* strains worldwide, we identified 20 distinct *AvrRvi6* alleles, of which only five are not recognized by Rvi6. Evolution analysis demonstrated that the emergence of these virulent alleles predates the domestication of apple, thus revealing that wild *Malus* species constitute a reservoir of virulence. Using AlphaFold structural modeling, we showed that AvrRvi6 belongs to an expanded family that adopts a MAX effector fold, originally described for cytoplasmic effectors of the blast fungus *Pyricularia oryzae*. Transient expression in *Nicotiana benthamiana* demonstrated that AvrRvi6 triggers allele-specific apoplastic immunity and validated three different molecular mechanisms (mutation, partial deletion and transposon insertion in the promoter) that *V. inaequalis* employs to circumvent *Rvi6* recognition. This study delivers the first characterization of a *V. inaequalis* avirulence factor, uncovers an unexpected apoplastic role for a MAX effector, and demonstrates how ancestral virulence diversity maintained in wild populations compromises resistance durability in apple.

## Introduction

The outcome of a plant–microbe interaction is determined at a molecular battleground where pathogens deploy virulence factors and plants deploy immune receptors to detect and restrict infection. Plant immune receptors survey both the extracellular space and the host cell interior, creating multiple layers of surveillance that pathogens must evade (Kourelis and van der Hoorn, 2018). This reciprocal pressure drives a continuous evolutionary arms race in which changes in pathogen virulence are countered by changes in host recognition. Deciphering this molecular dialogue is fundamental to understanding how plant resistance evolves, how pathogens overcome it, and ultimately how durable resistance can be achieved (Jones et al., 2024).

Apple is one of the most widely cultivated fruit crops worldwide (Muder et al., 2022). Despite the use of resistant cultivars, it is among the most heavily treated crops, mostly because of the extensive use of fungicides (often 12 to 25 applications per year) required to control apple scab disease, caused by the Ascomycota fungus *Venturia inaequalis* (Antal et al., 2024). This pathogen is present in nearly all apple-producing regions and, when poorly managed, can cause yield losses of 20–70% (Antal et al., 2024; Bowen et al., 2011). It is also capable of infecting the many wild *Malus* spp. found in the Northern Hemisphere (Bowen et al., 2011; Feurtey et al., 2020). After penetrating the cuticle, *V. inaequalis* follows an unusual infection strategy: it develops specialized subcuticular stromata and runner hyphae that remain confined between the cuticle and the epidermal cell wall, without forming intracellular feeding structures (Steiner and Oerke, 2024). This lifestyle is associated with a major reprogramming of carbohydrate metabolism and the secretion of hundreds of small proteins, known as effectors (Deng et al., 2017; Rocafort et al., 2022; Rocafort et al., 2023).

Effectors are key virulence determinants. They can be translocated into host cells or remain in the apoplast (Rocafort et al., 2020). In many plant–pathogen interactions, effectors manipulate host physiology to promote infection, for example by suppressing immune responses (Oliveira-Garcia et al., 2023) or modulating the microbiota (Snelders et al., 2022). While effector protein sequences are typically highly diverse, many of them can be grouped in effector families based on conserved folds (Derbyshire and Raffaele, 2023; Jones and Raffaele, 2025; Le Naour-Vernet et al., 2025; Rocafort et al., 2022; Seong and Krasileva, 2023). One of the best-studied families is the MAX (*Magnaporthe* Avr and ToxB-like) effector family, which comprises small proteins, typically less than 150 amino acids, characterized by a conserved six-stranded β-sandwich fold stabilized by disulfide bridges (de Guillen et al., 2015). Members of the MAX family are found in several plant-pathogenic fungi, but they are particularly expanded in *Pyricularia oryzae*, *Claviceps purpurea*, and *V. inaequalis* (de Guillen et al., 2015; Le Naour-Vernet et al., 2025). In *V. inaequalis*, MAX effectors represent the largest effector family, with 75 predicted members (Rocafort et al., 2022). Structural comparisons suggest that this family represents an evolutionary innovation allowing pathogens to diversify their effector repertoires while maintaining a robust and adaptable fold (Lahfa et al., 2024).

Although effectors are crucial for virulence, they were initially characterized as avirulence (AVR) proteins recognized by resistant cultivars. Indeed, plants have evolved immune receptors (encoded by resistance (*R*) genes) that directly or indirectly recognize fungal effectors (Cesari, 2018; Kourelis and van der Hoorn, 2018) and activate strong defense responses, often manifested as a hypersensitive response (HR) (Heath, 2000; Stergiopoulos and De Wit, 2009). *R* genes encode either membrane-bound receptors that typically detect apoplastic effectors, or intracellular Nucleotide-binding Leucine-rich Repeat (NLR) proteins that recognize cytoplasmic effectors (Kourelis and van der Hoorn, 2018). Several apple *R* genes conferring resistance to *V. inaequalis* have been genetically identified in apple germplasm (Bowen et al., 2011; Bus et al., 2011; Patocchi et al., 2020). To date, three apple *R* genes have been cloned and functionally validated: *Rvi4* (*Vr2c*, *Rvi15*), *Rvi6* (*HcrVf2*), and *Rvi12* (*Vb*) (Belfanti et al., 2004; Gaiii et al., 2010; Peil et al., 2023; Schouten et al., 2014). *Rvi4* encodes a TIR-NBS-LRR protein and likely recognizes a cytoplasmic effector (Schouten et al., 2014), whereas *Rvi6* and *Rvi12* encode membrane-bound receptors predicted to detect apoplastic effectors (Belfanti et al., 2004; Padmarasu et al., 2018), similar to the *Cf* immune receptors of tomato (Rivas and Thomas, 2005) and *RXEG1* of *Nicotiana benthamiana* (Wang et al., 2018). Thus, the apple–*V. inaequalis* pathosystem provides an opportunity to investigate how effectors operating in distinct cellular compartments are coupled to host immune recognition.

Among these resistance genes, *Rvi6* has been the most widely deployed *R* gene in commercial apple cultivars and was initially considered to provide durable resistance (Crosby et al., 1992). However, *V. inaequalis* populations capable of overcoming *Rvi6* were first reported in Germany in 1988 (Parisi et al., 1993), followed by reports in various European countries (Lemaire et al., 2016) and, more recently, the United States (Papp et al., 2020), but have not yet been reported in the Southern Hemisphere. *Rvi6* originated from the ornamental crabapple *Malus floribunda* and was introgressed into domesticated apple (*Malus × domestica*) (Dunemann et al., 2012). Population genetic studies revealed that *V. inaequalis* isolates virulent on *Rvi6* cultivars were genetically closer to isolates from *M. floribunda* than to those from susceptible *M. × domestica* (Gladieux et al., 2011; Guérin and Le Cam, 2004; Lemaire et al., 2016), until a secondary contact with gene flow between the two populations occurred in orchards on susceptible cultivars (Leroy et al., 2016). These studies suggest that virulent alleles pre-existed as standing variation in *V. inaequalis* populations outside orchards, which then expanded under agricultural selection pressure (Lemaire et al., 2016). These findings underscore the need to dissect gene-for-gene relationships in the apple–*V. inaequalis* interaction to understand the mechanisms of resistance circumvention, thereby improving resistance management and its durability.

Here, we identify *AvrRvi6* as the fungal determinant of *Rvi6* recognition through genomic mapping, and functional complementation. We show that AvrRvi6 belongs to the MAX effector family but, unexpectedly, functions in the apoplast rather than the host cytoplasm. Analysis of 122 *V. inaequalis* isolates reveals extensive natural variation at the *AvrRvi6* locus and identifies multiple molecular routes by which the pathogen escapes *Rvi6* recognition. Finally, population genomic analyses indicate that virulence-associated variation at AvrRvi6 predates apple domestication by thousands of years. Together, these findings reveal an unexpected extracellular function for a conserved fungal effector fold and demonstrate how ancient pathogen diversity can constrain the durability of contemporary plant immune resistance.

## Results

### 1. Genome-wide association mapping of the *AvrRvi6* locus

To identify the genomic region comprising *AvrRvi6*, we performed pooled-genome sequencing of two groups of *V. inaequalis* isolates that are either avirulent or virulent toward *Rvi6*-resistant apple cultivars. First, we pooled 24 and 27 mono-ascosporic progenies, from an *in vitro* cross between isolates 0301 and 1066, which are avirulent and virulent on *Rvi6*-resistant cultivars, respectively (Supplemental Table 1; (Bénaouf and Parisi, 2000; Broggini et al., 2007). Second, we leveraged the accumulation of recombination events in natural populations from a French orchard that had experienced a recent episode of intense secondary gene flow (Leroy et al., 2016). Among the 41 individuals isolated from *Rvi6*-resistant and non-*Rvi6* cultivars in this orchard (Supplemental Table 1), 22 and 19 isolates were avirulent and virulent, respectively, on an *Rvi6*-resistant cultivar (Supplemental Table 1). Sequencing reads from each pool were aligned to the reference genome of the avirulent EU-B04 isolate (Le Cam et al., 2019), followed by single nucleotide polymorphism (SNP) calling. *Fst* estimates were computed to identify genomic regions showing strong allele frequency differences between the avirulent and virulent pools from the natural populations or 0301 *×* 1066 progenies. The distribution of *Fst* estimates along the *V. inaequalis* genome highlighted scaffold UTG016 (2,111 kb) as the key region distinguishing avirulent and virulent progenies (Figure 1A). While *Fst* estimates from natural populations also highlighted scaffold UTG013 (Figure 1A), we focused on scaffold UTG016 because it was the only one sharing a clear signal in both natural populations and progenies. The analysis of *Fst* estimates from natural populations further narrowed the candidate region on scaffold UTG016 to 95 kb (positions 1,679,308–1,773,969) (Figure 1B). This region corresponds to a GC-equilibrated isochore flanked by AT-rich regions (Figure 1C) and contains 25 predicted genes, most of which exhibit transcriptional support from *in planta* RNA-seq expression data (Figure 1D; Supplemental Table 2). Because most *AVR* genes typically encode small secreted proteins (SSPs) (Rocafort et al., 2020), we hypothesized that *AvrRvi6* would also encode such an effector, *i.e.* a peptide of less than 300 amino acids, cysteine-rich and with a signal peptide. Among the 25 predicted genes at this locus, only two encode proteins with a predicted signal peptide. Furthermore, only one of them, *g11287*, shows strong expression at 48 hours post-inoculation based on RNA-seq data from apple leaves infected with *V. inaequalis* isolate EU-B04 (Figure 1E; Supplemental Table 2). In the isolate MNH120, *g11287* shows a strong induction in inoculated condition with a peak of expression at 3- and 5-days post inoculation (Figure 1F, (Rocafort et al., 2022)) Thus, *g11287* was identified as the single candidate gene for *AvrRiv6*.

**Figure 1:**
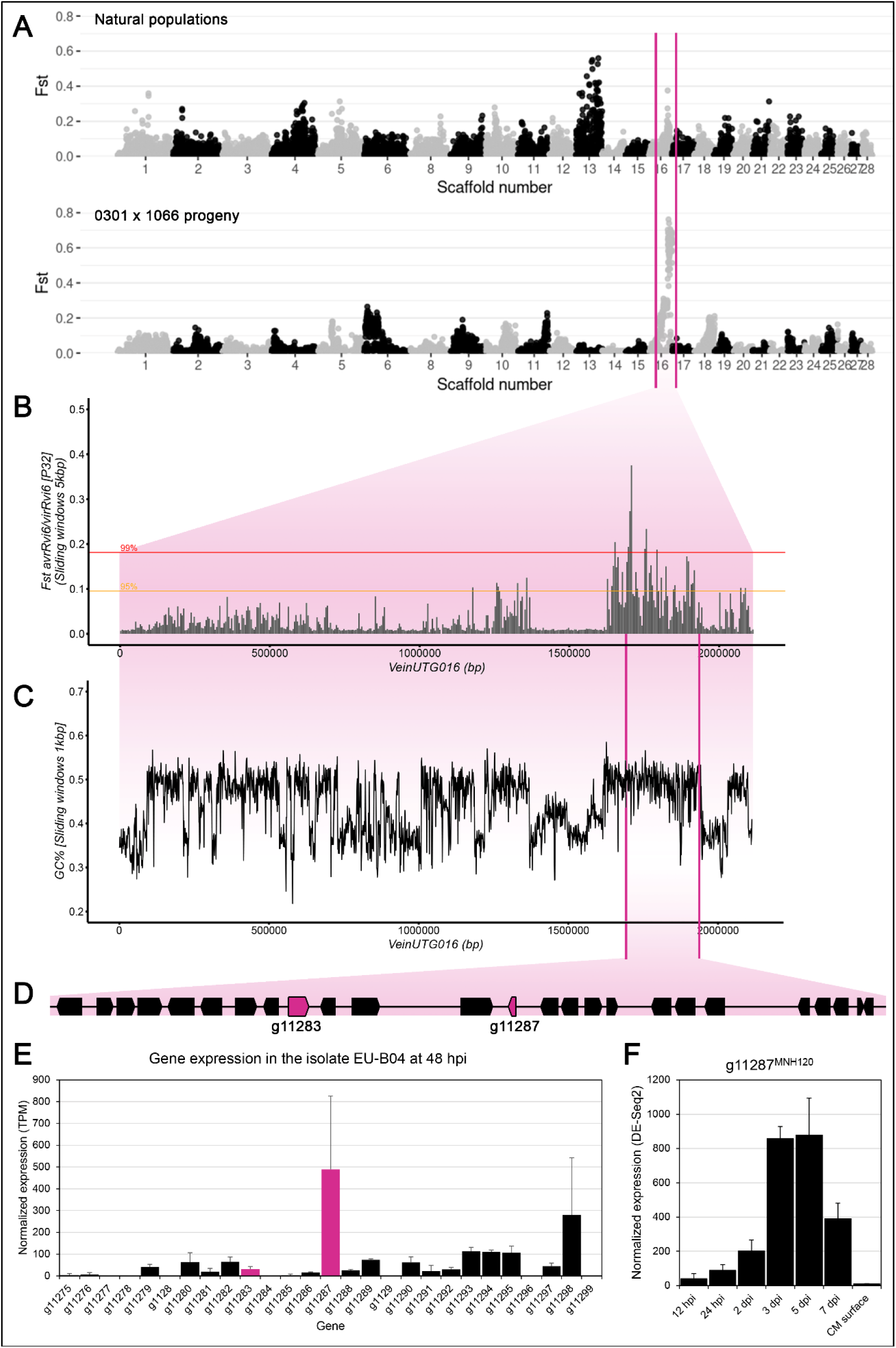
Genome-wide association mapping to identify *AvrRvi6*. **A.** *Fst* estimates of SNPs were determined between wild avirulent and virulent isolates (natural populations), and between avirulent and virulent progenies from a cross between 0301 and 1066 isolates. **B.** *Fst* estimates of SNPs between avirulent and virulent natural populations for the VeinUTG016 scaffold. **C.** GC content across the VeinUTG016 scaffold. **D.** Schematic representation of genes and their orientation on scaffold UTG016 (positions 1,679,308– 1,773,969), secreted proteins g11283 and g11287 are highlighted in magenta. **E.** Expression for each candidate genes are shown as bar graphs, data are from isolate EU-B04 inoculated on apple cultivar Gala at 48 hours post-inoculation (hpi) with three independent biological replicates. **F.** Gene expression of *g11287* in the isolate MNH120, data from Rocafort et al., 2022. TPM, Transcripts Per Million; Fst, Fixation Index; dpi, day post inoculation; CM, cellophane membrane.

To confirm that *g11287* is *AvrRvi6*, we introduced the EU-B04 allele under the control of its native promoter in the virulent isolate 2828 (Supplemental Figure 1). Two independent transformants failed to produce symptoms on resistant *Rvi6* cultivars J45 and Priscilla while retaining the ability to colonize the susceptible cultivars Gala and Golden Delicious (Figure 2; Supplemental Figure 2). These results demonstrate that transgenic expression of the *g11287* gene from EU-B04 in isolate 2828 triggers *Rvi6*-mediated immunity, thereby confirming that the SSP encoded by *g11287* is AvrRvi6.

**Figure 2:**
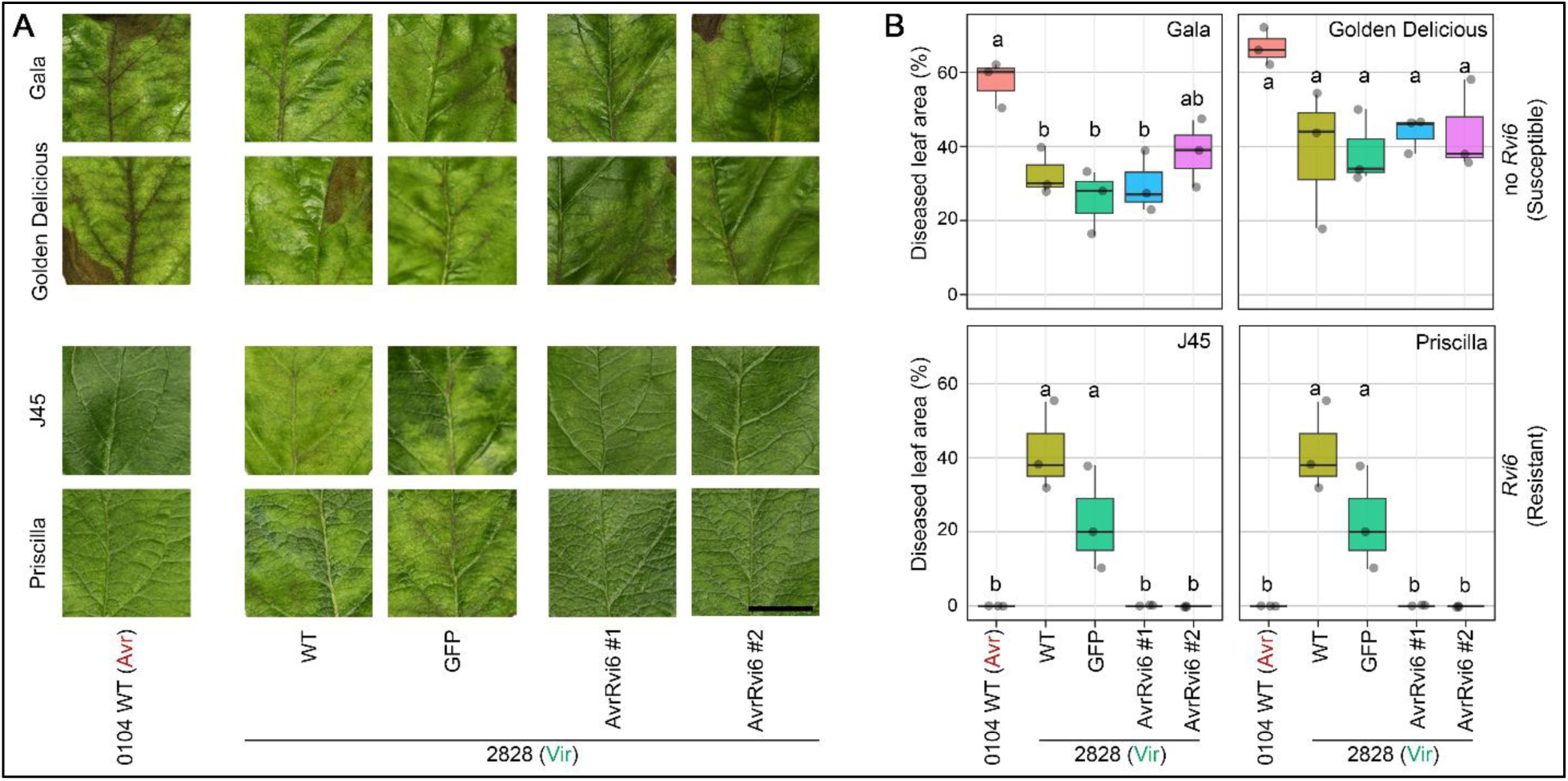
Functional validation of *g11287* by complementation of the virulent isolate 2828. Wild-type (WT) isolates 0104, 2828, as well as the corresponding transformants expressing either green fluorescent protein (GFP) or *pAvrRvi6::AvrRvi6^EU-B04^*(two independent transformants AvrRvi6 #1 and AvrRvi6 #2), were inoculated onto young grafted apple trees. The Gala and Golden Delicious cultivars lack the *Rvi6* resistance gene, whereas J45 and Priscilla carry *Rvi6*. **A.** Representative symptoms at 21 days post-inoculation on the f0 leaf (the last fully expanded leaf at the time of inoculation). Scale bar = 1 cm. **B.** Average diseased leaf area (%) at 21 days post-inoculation from five leaves per plant, plotted for a minimum of three independent plants. Letters indicate significance groups as determined by one-way ANOVA followed by Tukey’s post hoc test (*P* ≤ 0.05).

### 2. AvrRvi6 belongs to the MAX effector family, which is expanded in *Venturia inaequalis*

The *AvrRvi6* gene is 335 base pairs (bp) long, contains a single 53 bp intron, and encodes a protein with typical features of a fungal effector. More specifically, the AvrRvi6 protein is small (93 amino acids), cysteine-rich (contains six cysteine residues), carries an N-terminal signal peptide for secretion (Figure 3A), and shows no sequence homology to other known proteins. As no function or known conserved domain could be predicted, the AvrRvi6^EU-B04^ protein structure was modeled using AlphaFold2 (Mirdita et al., 2022) to get further insights and the highest ranking model was chosen based on its confidence scores. The AvrRvi6^EU-B04^ structure was predicted with very high confidence (pLDDT score of 95.5 and pTM score of 0.804). The AvrRvi6^EU-B04^ model showed a six-stranded β-sandwich fold: β1, β2 and β6 form one β-sheet, while β3, β4 and β5 form a second (Figure 3B), which is typical of MAX effectors (de Guillen et al., 2015). The predicted structure is stabilized by three disulfide bonds: SS1: β1 Cys3 – α1 Cys 43; SS2: β1 Cys4 – α3 Cys 74; SS3: β4 Cys32 – 21 Cys 56 (Figure 3C). Disulfide bond 1, located between the conserved cysteines in the beginning of β1 and just before β5, is also typical of MAX effectors (de Guillen et al., 2015). The closest structural homolog of AvrRvi6 in the Protein Data Bank (PDB) is MoToxB (PDB ID: 6RJ5), which belongs to the MAX structural family, with a significant Z score of 5 (Lahfa et al., 2024) (Figure 3D). Taken together, protein structural modeling confirms that AvrRvi6 belongs to the MAX effectors family.

**Figure 3:**
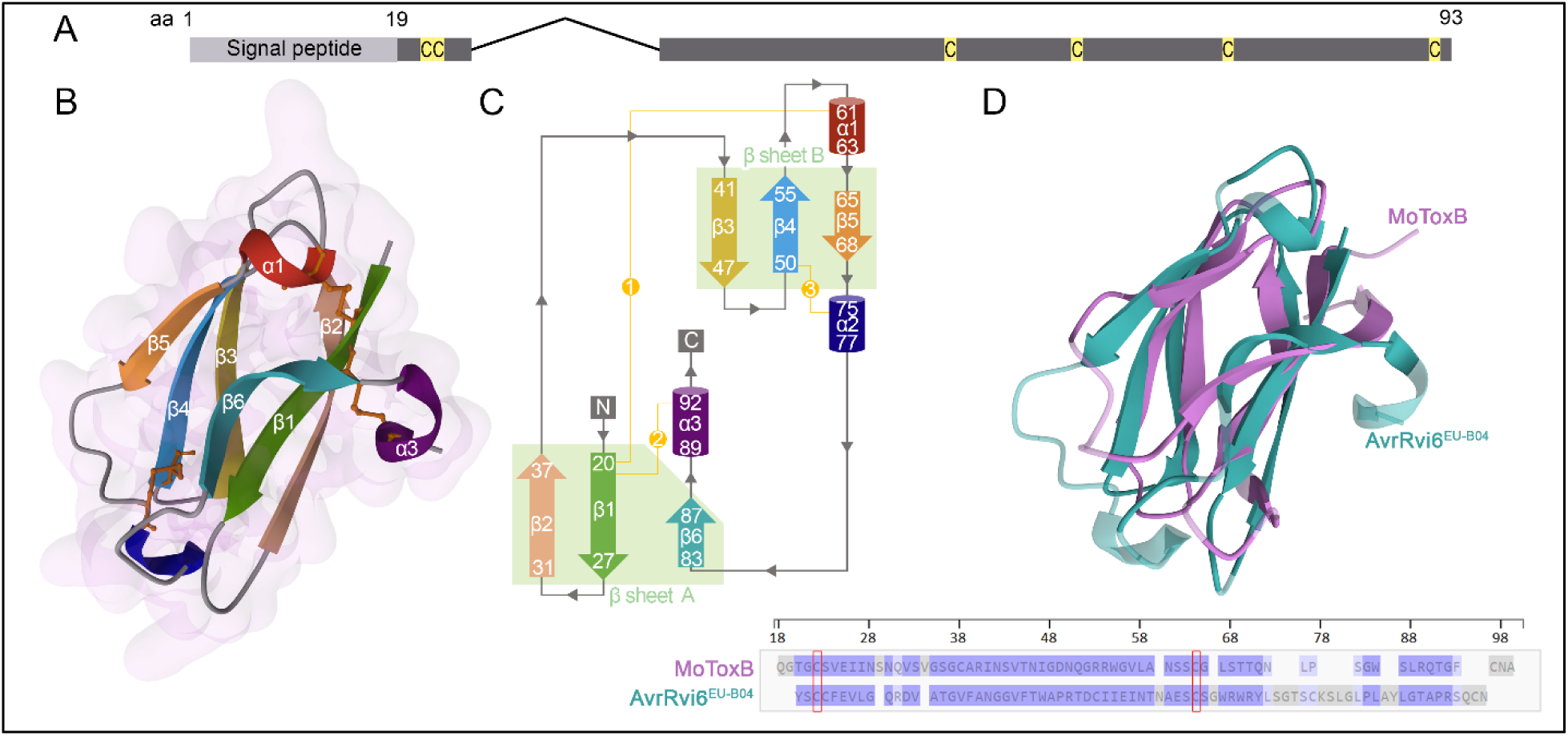
AvrRvi6 is a MAX effector. **A.** Schematic representation of the AvrRvi6 protein. The letter “C” marks the positions of cysteine residues. **B.** Cartoon representation of the AvrRvi6^EU-B04^ AlphaFold2-predicted structure, colored in a rainbow gradient, with disulfide bonds shown as orange sticks and the surface view in light pink. Predicted Local Distance Difference Test (pLDDT) score = 95.5; predicted Template Modeling score (pTM) = 0.804. **C.** Topology diagram of AvrRvi6 drawn by PDBsum and colored as in panel A. The two β-sheets are labeled A and B, with strands indicated by arrows and three disulfide bonds shown as yellow dotted lines. **D.** 3D structural alignment of AvrRvi6^EU-B04^ with MoToxB (PDB ID: 6RJ5) using PDBalign (Template Modeling Score (TM-score) = 0.56). Highly conserved cysteine residues among MAX effectors are outlined in red.

The MAX effector gene family has previously been reported to be massively expanded in the genome of *V. inaequalis* isolate MNH120 (Rocafort et al., 2022). In addition to *AvrRvi6*, 112 paralogous (*AvrRvi6*-like) sequences were identified in the EU-B04 genome, of which 75 were predicted to encode full-length proteins with a signal peptide and the six above-mentioned cysteine residues. Similar to *AvrRvi6*, 50 of these 75 *AvrRvi6*-like sequences were predicted to contain one intron, while 18 contained two introns and eight contained none (Supplemental Table 3). Alignment of the 75 AvrRvi6-like proteins with AvrRvi6 revealed a high level of sequence conservation across the signal peptide. However, beyond the signal peptide, sequence conservation was restricted to the six cysteine residues, as well as a limited number of other amino acid residues anticipated to be required for structural conservation of the MAX fold (Supplemental Figure 3). Percentage sequence identity to AvrRvi6 ranged from 27.6% to 50.5% (Supplemental Table 4).

### 3. AvrRvi6 is localized and detected in the apoplast

Because *Rvi6* encodes a receptor-like protein (RLP) with a predicted extracellular leucine-rich repeat (LRR) domain (Belfanti et al., 2004), we hypothesized that AvrRvi6 detection by Rvi6 occurs in the apoplast. To address this hypothesis, we transiently expressed AvrRvi6 in *N. benthamiana* leaves with or without a signal peptide. To ensure proper secretion into the apoplast, the native signal peptide (amino acids 1–19) was replaced with the Pathogenesis-Related 1α (PR1α) signal peptide from *Nicotiana tabacum*. When expressed individually, neither PR1α^SP^AvrRvi6^ΔSP^ nor Rvi6 induced an immune response (Figure 4A). However, co-expression of PR1α^SP^AvrRvi6^ΔSP^ with Rvi6 triggered a strong hypersensitive response (HR) four days after agroinfiltration, similar to the positive control combining the Cf-4 RLP from tomato (Thomas et al., 1997) and its corresponding apoplastic effector from the tomato leaf mold fungus *Fulvia fulva* (syn. *Cladosporium fulvum*), PR1α^SP^Avr4^ΔSP^ (Joosten et al., 1994). This recognition was specific, as neither Cf-4 co-expressed with PR1α^SP^AvrRvi6^ΔSP^nor Rvi6 co-expressed with PR1α^SP^Avr4^ΔSP^ triggered an HR (Figure 4A). In contrast, co-expression of AvrRvi6^ΔSP^, which restricts AvrRvi6 to the cytoplasm of the plant cell, and Rvi6 did not induce an HR, confirming that AvrRvi6 recognition requires secretion into the apoplast (Figure 4B).

**Figure 4:**
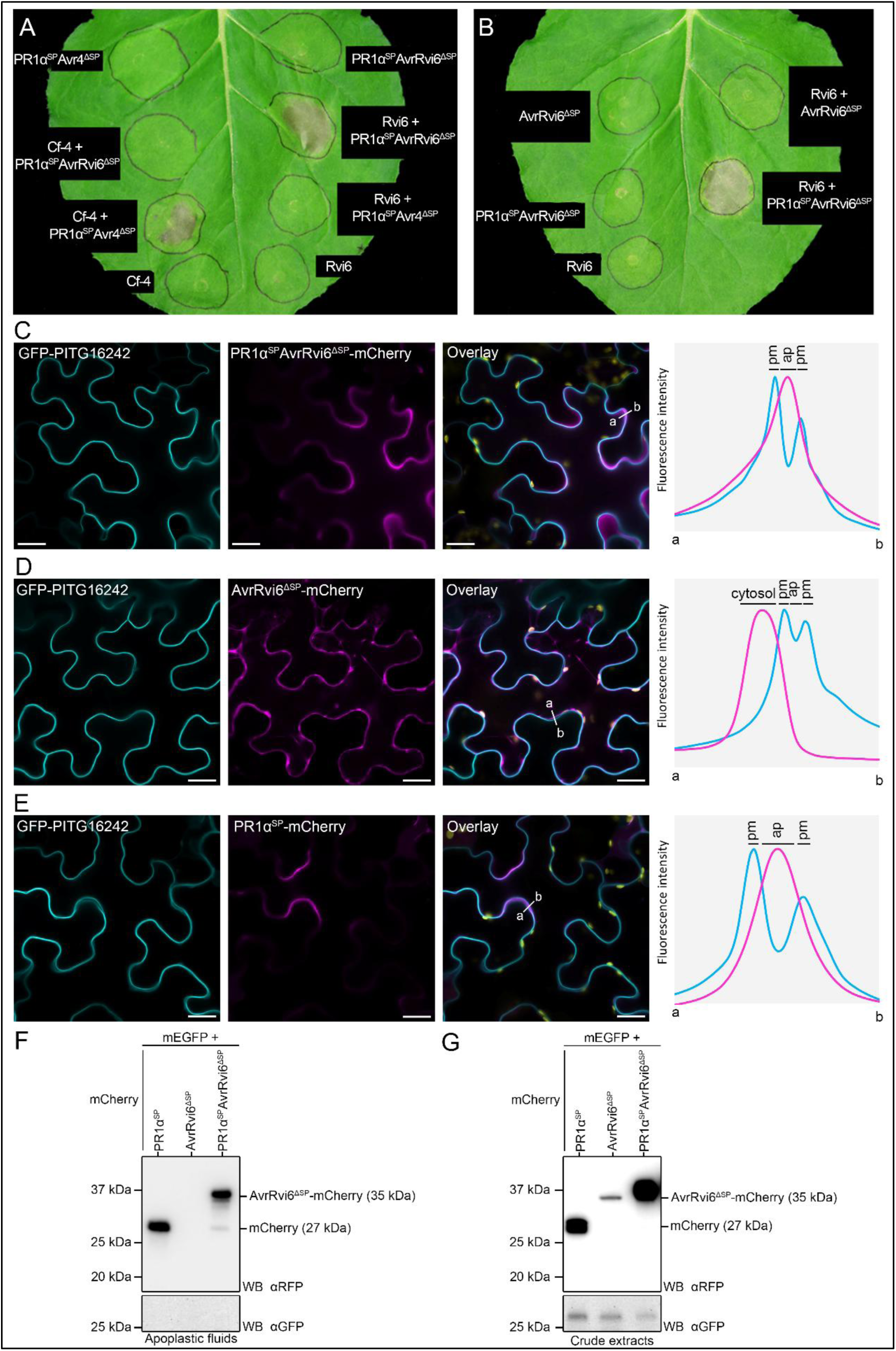
AvrRvi6 is localized in the *Nicotiana benthamiana* apoplast. **A.** *Agrobacterium tumefaciens* carrying constructs expressing PR1α^SP^Avr4^ΔSP^, Cf-4, PR1α^SP^AvrRvi6^ΔSP^, and Rvi6 were infiltrated in *N. benthamiana* alone or in combination. Images were taken 4 days post-infiltration, with pictures showing representative results from three independent biological replicates. **B.** *A. tumefaciens* carrying constructs expressing AvrRvi6^ΔSP^, PR1α^SP^AvrRvi6^ΔSP^, and Rvi6 were infiltrated in *N. benthamiana* alone or in combination. The picture, taken 4 days post-infiltration, shows representative results from three independent biological replicates. **C** – **E.** PR1α^SP^AvrRvi6^ΔSP^**-**mCherry (**C**), AvrRvi6^ΔSP^**-**mCherry (**D**) or PR1α^SP^**-**mCherry (**E**) were co-expressed with the plasma membrane marker GFP-PITG16242 *via* agroinfiltration in *N. benthamiana* leaves. Live-cell imaging was performed with a laser-scanning confocal microscope 3 days after infiltration. Images are single optical sections. The overlay panel combines GFP, mCherry and chlorophyll (yellow) channels. The right panels show relative fluorescence intensity plots of the GFP (cyan) and the mCherry (magenta) signals along the line from a to b depicted in the corresponding overlay panels. pm = plasma membrane, ap = apoplast. Bar = 20 µm. **F** – **G** Western Blot (WB) analysis on apoplastic wash fluids (**F**) or whole protein extracts (**G**) from *N. benthamiana* leaves expressing free mEGFP and PR1α^SP^AvrRvi6^ΔSP^-mCherry, AvrRvi6^ΔSP^-mCherry or PR1α^SP^-mCherry. Membranes were first hybridized with anti-RFP antibody, then stripped and re-hybridized with anti-GFP antibody.

To confirm the localization of PR1α^SP^AvrRvi6^ΔSP^, we fused the low acid sensitivity fluorescent protein mCherry to its C-terminal end and transiently expressed the construct in *N. benthamiana.* Laser-scanning confocal microscopy was then used to localize PR1α^SP^AvrRvi6^ΔSP^-mCherry co-expressed with the plasma membrane-localized RXLR effector GFP-PITG16242 (Petre et al., 2021) and the silencing suppressor P19 in epidermal cells. We observed PR1α^SP^AvrRvi6^ΔSP^-mCherry fluorescence signal at the cell periphery; however, it did not overlap with the plasma membrane-localized GFP signal (Figure 4C). Fluorescence signal analysis shows that PR1α^SP^AvrRvi6^ΔSP^-mCherry is found solely in the intercellular spaces between plasma membranes, with a very similar localization profile to that of the mCherry fused to PR1α^SP^ alone (PR1α^SP^-mCherry) (Figure 4C & 4D). In contrast, the control AvrRvi6^ΔSP^-mCherry, without a signal peptide, remained in the cytoplasm (Figure 4E). This indicates that PR1α^SP^AvrRvi6^ΔSP^-mCherry is, as expected, secreted into the apoplast and accumulates there without being re-internalized into the plant cells. The integrity of the tagged proteins was tested by western blot analysis on apoplastic wash fluids (Figure 4F) and crude protein extracts (Figure 4G). Only PR1α^SP^-mCherry and PR1α^SP^AvrRvi6^ΔSP^-mCherry were detected in apoplastic wash fluids, whereas the crude extracts showed expression of all three mCherry protein constructs, as well as free cytoplasmic mEGFP (Figure 4F and G). While cleavage of the mCherry tag was observed for PR1α^SP^AvrRvi6^ΔSP^-mCherry in apoplastic wash fluids, the majority of the accumulated protein was detected at the expected full-length size, demonstrating that the signal in Figure 4C mainly represents AvrRvi6 localization. Taken together, the HR assay and cellular imaging establish AvrRvi6 as the first characterized MAX effector to trigger immunity via a membrane-bound receptor.

### 3. Allelic diversity of AvrRvi6 in *Venturia inaequalis* populations

To better characterize the *AvrRvi6* diversity in *V. inaequalis* populations and identify molecular mechanisms that lead to *Rvi6* circumvention, we sequenced the *AvrRvi6* locus in 122 fungal isolates (Supplemental Table 5) and identified 20 unique alleles distributed worldwide (Figure 5), which mainly diverge by either deletion or non-synonymous mutations (Supplemental Figure 4). Among the 122 isolates, we phenotyped 74 of them for their ability to infect *Rvi6*-resistant apple trees and classified them as virulent or avirulent in the presence of *Rvi6* (Figure 5B, Supplemental Table 5). We also inferred a virulent phenotype for 14 additional strains collected from *Rvi6* cultivars (Figure 5B, Supplemental Table 5).

**Figure 5:**
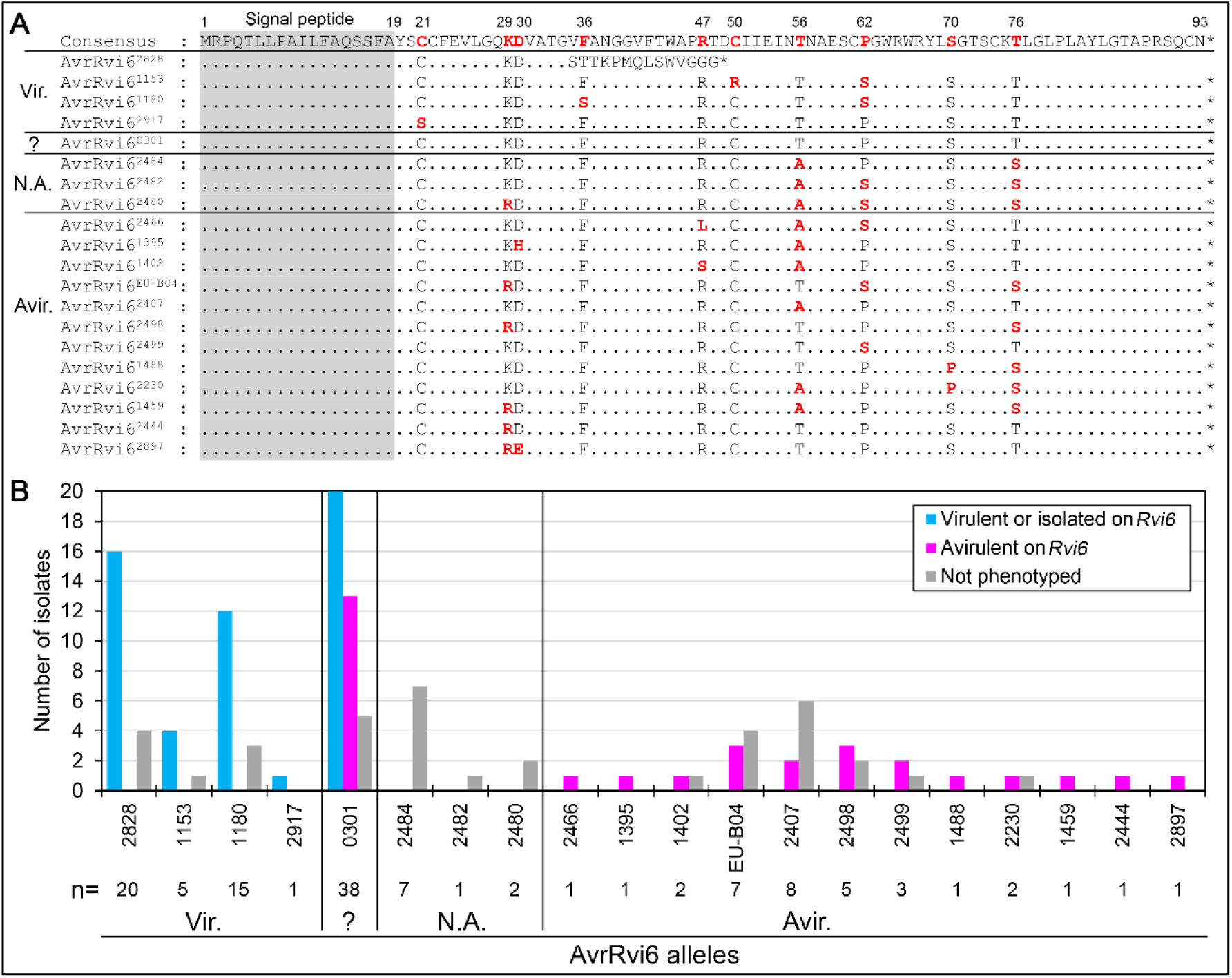
Allelic diversity of AvrRvi6 in *Venturia inaequalis* populations. **A.** The AvrRvi6 protein sequence alignment is shown for each of the 20 AvrRvi6 alleles; polymorphic residues are in red, and the signal peptide is highlighted in grey. The consensus sequence is shown at the top. **B.** Bar graph showing the number of fungal isolates carrying the different AvrRvi6 alleles and their phenotype on *Rvi6*-resistant trees. The total number of isolates for each allele is indicated below the bar graph (n=). Vir. = virulent,= variable, N.A. = non-available, Avir. = avirulent.

We found only four alleles (AvrRvi6^2828^, AvrRvi6^1153^, AvrRvi6^1180^, and AvrRvi6^2917^) that were consistently associated with virulent isolates, indicating variants that circumvented recognition by *Rvi6*. Compared to the most common avirulent allele AvrRvi6^0301^, amino acid polymorphisms associated with virulence were C21S (AvrRvi6^2917^), F36S (AvrRvi6^1180^), C50R (AvrRvi6^1153^) and a partial deletion causing a frame shift leading to an early stop codon (AvrRvi6^2828^), which also contained the likely disruptive mutation F36T and undetermined R47G. While the mutation of cysteines is expected to alter AvrRvi6 stability as reported for Avr4 (Van den Burg et al., 2003), the phenylalanine F36 residue might be important for the direct or indirect recognition by Rvi6. The AvrRvi6^1180^ allele also contains a P62S mutation, which is found in the majority of avirulent alleles and therefore is unlikely to play a role in evasion of recognition (Figure 5A).

Intriguingly, the AvrRvi6^0301^ allele, which is the most abundant in our sampling, was present in both virulent and avirulent isolates (Figure 5A). Promoter analysis of avirulent and virulent isolates carrying the AvrRvi6^0301^ allele revealed that avirulent isolates have a promoter similar to that of AvrRvi6^EU*-*B04^, whereas virulent isolates show a different promoter sequence (Supplemental Figure 5A). Indeed, we identified in isolate 1066 a 288 bp-long insertion with features of Miniature Inverted-repeat Transposable Elements (MITEs) (Tossolini et al., 2025) 15 nucleotides upstream of the ATG of *AvrRvi6* (Supplemental Figure 5B). We hypothesized that virulent isolates with this transposon insertion might escape recognition due to lack of gene expression. RT-qPCR analysis showed high *AvrRvi6* expression in the avirulent isolate EU-B04 during infection, whereas *AvrRvi6* transcripts were barely detectable in the virulent isolate 1066, confirming that escape from *Rvi6* resistance was due to lack of expression (Supplemental Figure 5C).

Allelic diversity was also detected among avirulent isolates with fourteen different alleles. Apart from *AvrRvi6^0301^*, *AvrRvi6^2466^*, *AvrRvi6^2480^*, *AvrRvi6^1395^*, *AvrRvi6^2476^*, *AvrRvi6^EU-B04^*, *AvrRvi6^2407^*, *AvrRvi6^2498^*, *AvrRvi6^2499^*, *AvrRvi6^1488^*, *AvrRvi6^2230^*, *AvrRvi6^1459^*, *AvrRvi6^2444^* and *AvrRvi6^2897^* were always found in avirulent isolates, indicating allelic variants that remain recognized by *Rvi6* (Figure 5B). Alleles *AvrRvi6^2480^*, *AvrRvi6^2484^* and *AvrRvi6^2482^* were exclusively found in isolates sampled from wild *Malus* species (*M. sieversii*, *M. sylvestris* or *M. orientalis*), and their virulence could not be assessed on *Rvi6* cultivars because they are non-pathogenic on susceptible *M. × domestica* (Feurtey et al., 2020). However, mutations in these alleles are shared among other *AVR* alleles and thus are expected to lead to recognition (Figure 5A). Population-wide analysis thus highlights substantial allelic diversity in both the promoter and coding sequence of the *AvrRvi6* gene, underscoring at least three different mechanisms that led *V. inaequalis* to evade *Rvi6*-triggered immunity.

### 4. Natural polymorphisms in *AvrRvi6* mediate evasion of *Rvi6* immunity

Because transient co-expression of Rvi6 and AvrRvi6 in *N. benthamiana* produced a strong HR, we used this system to investigate the impact of natural polymorphisms identified in *AvrRvi6* alleles on *Rvi6*-mediated immunity. We then co-expressed each AvrRvi6 allele with either Rvi6 or a control receptor (Cf-4), which is not involved in AvrRvi6-triggered immunity. HR intensity was quantified using red-light imaging (Landeo Villanueva et al., 2021) and compared to the HR induced by Cf-4 and Avr4. The four alleles associated with virulent isolates (AvrRvi6^2828^, AvrRvi6^2199^, AvrRvi6^1180^, and AvrRvi6^2917^) exhibited similar signal to the negative control Avr4 as they did not trigger an Rvi6-mediated HR, confirming that polymorphisms in these alleles impair Rvi6 recognition (Figure 6A and 6B). Certain replicates showed an HR, even with Avr4, suggesting slight Rvi6 self-activation when over-expressed (Chen et al., 2025) (Figure 6). The allele *AvrRvi6^0301^*, found in both virulent and avirulent strains, did induce an HR, validating that *Rvi6* virulence in the strains with allele *AvrRvi6^0301^*is caused by the promoter disruption (Figure 6). For the three alleles AvrRvi6^2484^, AvrRvi6^2482^ and AvrRvi6^2480^ that were found in isolates non-pathogenic on *M. × domestica*, we observed a strong HR suggesting that the isolates carrying these alleles would be avirulent on *Rvi6*-carrying trees (Figure 6A and B). This result is coherent with the shared polymorphism (T56A, P62S and T76S) with other avirulent alleles. Among the thirteen alleles associated with avirulent strains, the nine alleles that were tested (AvrRvi6^0301^, AvrRvi6^2466^, AvrRvi6^1395^, AvrRvi6^1402^, AvrRvi6^EU-B04^, AvrRvi6^2483^, AvrRvi6^2499^, AvrRvi6^1488^, and AvrRvi6^2230^) specifically trigger Rvi6-mediated HR with different intensity (Figure 6A and B). For the alleles AvrRvi6^2466^ and AvrRvi6^1488^ the weaker signal and variability is not significantly different than the negative control Avr4-Rvi6. Compared to the Avr4-Cf-4 recognition, the HR triggered by Rvi6 proved more variable as showed by the much larger standard deviation for most *AVR* alleles. To exclude a technical issue with transient expression, protein accumulation of AvrRvi6 alleles was tested by western blot analysis. All except the AvrRvi6^2828^ allele shown to be expressed at similar levels (Figure 6C), suggesting that variation in recognition of a given allele by Rvi6 is intrinsic to their interaction.. In conclusion, the *N. benthamiana* system recapitulates the phenotypes observed in *V. inaequalis* isolates on *Rvi6* apple trees and highlights that the non-synonymous mutations C21S, F36S, and C50R enable escape from *Rvi6*-mediated immunity.

**Figure 6:**
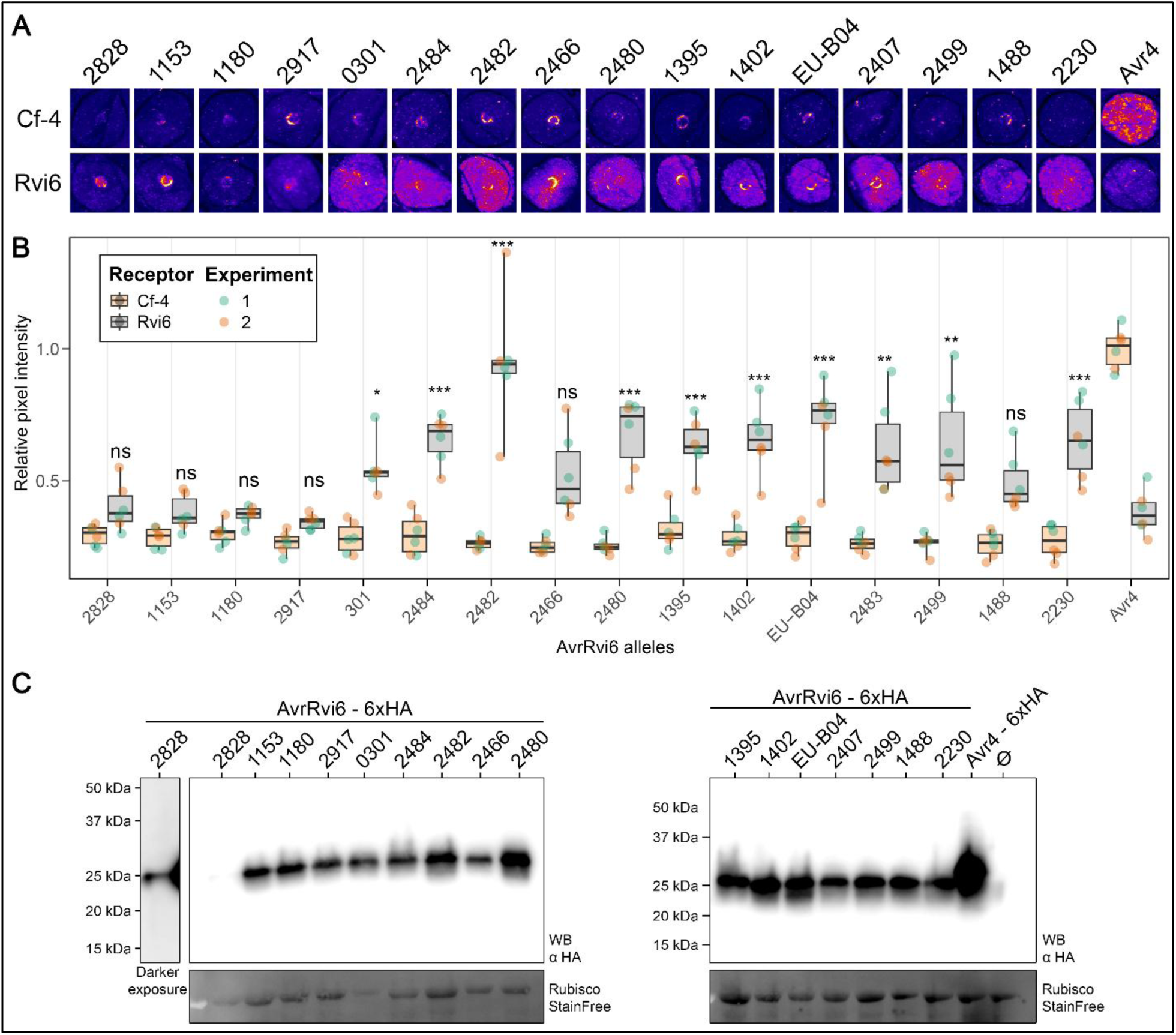
Allele-dependent activation of Rvi6-mediated immunity by AvrRvi6 in *Nicotiana benthamiana*. *Agrobacterium tumefaciens* carrying constructs expressing AvrRvi6 alleles or Avr4 fused to 6 × hemagglutinin (HA) epitope were co-infiltrated with Rvi6 or the control receptor Cf-4 into *N. benthamiana* leaves. Leaves were harvested 4 days post-infiltration, and hypersensitive response (HR) intensity was quantified using red-light imaging. **v.** Pictures of infiltrated spot imaged using Cy3 setting on Chemidoc® apparatus and colorized with Fire LUT settings on FIJI softwar. **B.** Quantification of pixel intensity for each allele was compared to the HR induced by Cf-4 and Avr4, which was co-infiltrated on each leaf. Boxplots represent data from two independent biological experiments, each comprising three leaves. Statistical comparisons among alleles were performed using an ANOVA followed by a Dunnet test with Avr4 as the reference, for Cf-4 and Rvi6 separately. For Cf-4, only Avr4 differs from the other alleles. For *Rvi6*, the statistical difference between each allele and Avr4 is indicated by ns = not significant; ** = *p* < 0.01; *** = *p* < 0.001. **C.** Western Blot (WB) analysis of AvrRvi6 alleles and Avr4.

### 5. Evolutionary history of AvrRvi6 allelic variation

To better understand the evolutionary origin of the polymorphisms observed in *AvrRvi6*, we combined sequence comparisons with network analysis. All non-synonymous polymorphisms identified in *AvrRvi6* can be generated by single-nucleotide substitutions, including the mutations associated with virulent alleles (Figure 7A). The topology of the network is consistent with *AvrRvi6^2917^*having originated from *AvrRvi6^0301^*, whereas *AvrRvi6^1153^*and *AvrRvi6^1180^* appear to have independently derived from an *AvrRvi6^2499^* background through additional mutations (Figure 7B). Several avirulent alleles are also closely related and differ by single substitutions, such as *AvrRvi6^2407^* and *AvrRvi6^1395^* or *AvrRvi6^2444^* and *AvrRvi6^2897^*. However, most alleles carry distinct combinations of polymorphisms, making their evolutionary relationships difficult to resolve from the network alone (Figure 7B). The four substitutions found exclusively in single alleles (D30H, D30E, R47L, and R47S) are therefore likely to represent more recent mutations, whereas substitutions shared among multiple alleles are consistent with older, ancestral polymorphisms.

**Figure 7:**
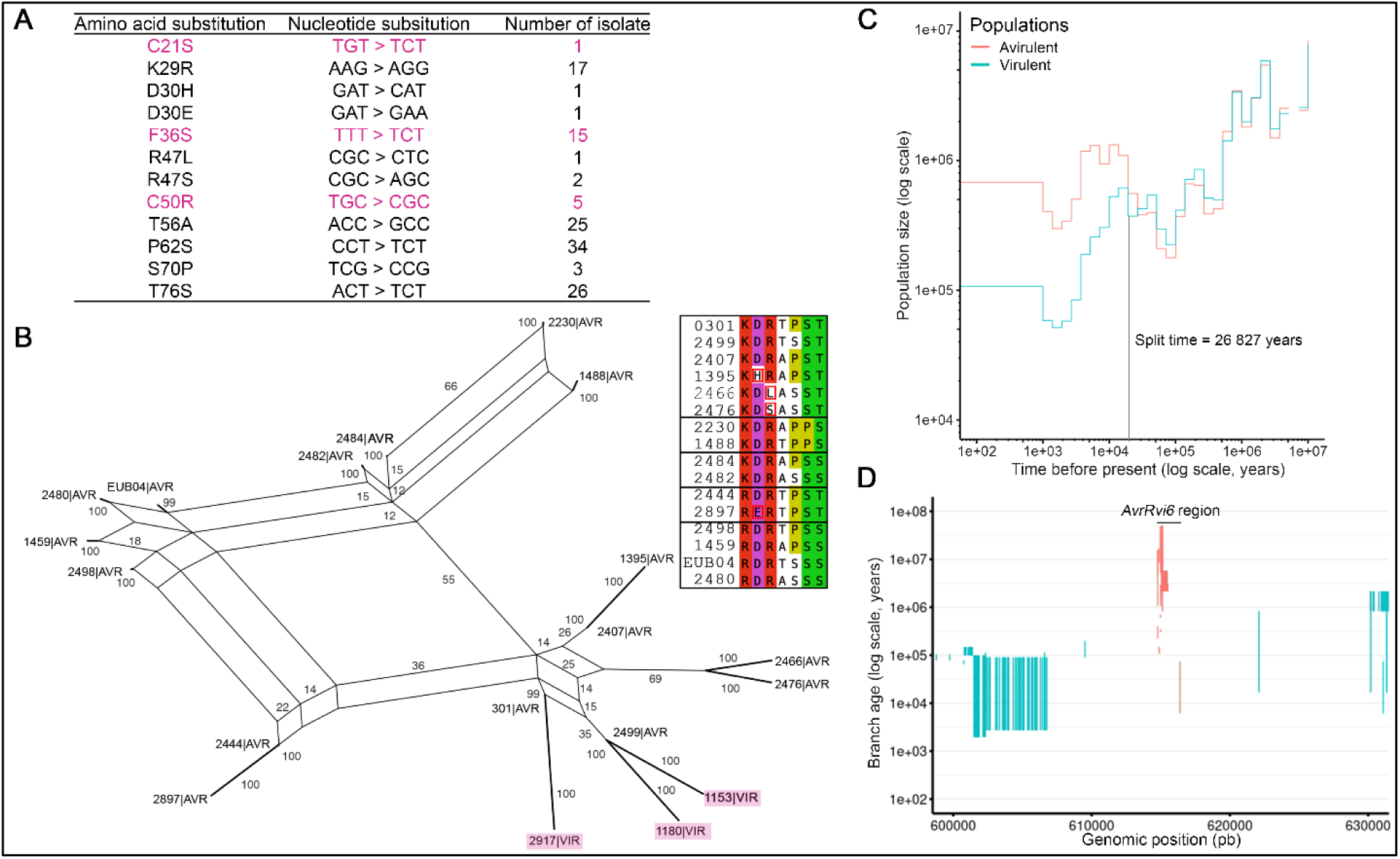
Evolutionary analyses of *AvrRvi6* alleles in *Venturia inaequalis*. **A.** Nucleotide and corresponding amino acid substitutions found in AvrRvi6 alleles. Virulent mutations are in magenta **B.** Amino acid-based network of AvrRvi6 alleles built with SplitsTree App 6.0.0 (Huson and Bryant 2024). Numbers indicate splits networks bootstrap. Polymorphic residues are showed on the right. Black boxes correspond to sub-networks and red boxes highlight unique substitutions. **C.** Demographic analysis of the *Rvi6-*avirulent (in red) and *Rvi6-* virulent (in blue) populations. **D.** Age of branches carrying a mutation on a genomic portion containing the scaffold VeinUTG016. The *AvrRvi6* region (1kb around) is indicated in red. This analysis was done on the *Rvi6*-avirulent population.

To investigate the evolutionary history of virulent and avirulent *V. inaequalis* populations, we analyzed the polymorphisms observed these populations from 38 isolates with genomic data (Supplemental Table 5). Based on 1,062,609 autosomal SNPs, we estimated population sizes and divergence times (Figure 7C). The two populations diverged *ca.* 27,000 years ago, after which the avirulent population underwent a strong expansion, while the virulent population experienced a decline. Current population sizes were estimated to be approximately 700,000 individuals for the avirulent population and 110,000 individuals for the virulent population. Both populations underwent a demographic bottleneck between 3,000 and 1,400 years ago. Positive selection analyses did not reveal any clear signature of adaptive processes in the *AvrRvi6* region of the virulent population (Supplemental Figure 6A-B). Finally, we estimated the ages of branches where mutations occurred. Figure 7D shows the lower and upper age estimates for each mutation within a ±1 kb window around the *AvrRvi6* gene (see also Supplemental Figure 6). Around *AvrRvi6*, lower age estimates ranged from 6,081 to 15,201,600 years, while upper age estimates ranged from 74,245 to 50,341,000 years. These results indicate that the polymorphisms observed in the *AvrRvi6* region reflect very ancient mutation events rather than recent adaptation to domesticated apple trees.

## Discussion

### *AvrRvi6*: the first avirulence gene identified in *Venturia inaequalis*

In this study, we identified and characterized *AvrRvi6*, the first avirulence gene discovered in *Venturia inaequalis*, the causal agent of apple scab. This discovery represents a milestone in the molecular understanding of the *V. inaequalis*-apple pathosystem because it closes a longstanding knowledge gap: although the corresponding resistance gene *Rvi6* was cloned and functionally validated over two decades ago (Belfanti et al., 2004), the corresponding *AVR* gene had remained elusive until now. Our study revealed three molecular mechanisms to overcome *Rvi6*-mediated resistance: non-synonymous mutations (especially of cysteine residues), partial deletion, and transposon-insertion in the promoter that leads to lack of expression. Each of these mechanisms have been reported in other plant-pathogenic fungi (Wang et al., 2022). Identifying the *AvrRvi6* gene was crucial for the monitoring of pathogen populations (Le Cam et al., 2021), and our study now enables the detection of virulent alleles, guiding growers in the deployment of *Rvi6* resistant cultivars (Le Cam et al., 2021).

### The emergence of virulence predates agriculture

Our population genomic analyses provide key insights into the evolutionary history of virulent and avirulent *V. inaequalis* populations. Coalescent-based modeling revealed that the divergence between *Rvi6*-virulent and avirulent populations occurred approximately 27,000 years ago, a result highly consistent with earlier estimates based on microsatellite data (∼23,700 years; (Lemaire et al., 2016). In many crop pathosystems, modern agriculture has driven the emergence of virulent pathogen strains. In contrast, our finding indicates that wild apple relatives present in the northern hemisphere, rather than modern agriculture, shaped this early diversification of *AvrRvi6* alleles. Consistent with this, the first report of *Rvi6* resistance breakdown in the United Kingdom occurred in 1989 on a wild *Malus floribunda* tree in a Kent garden (Roberts and Crute, 1994), well before *Rvi6* cultivars were deployed in English orchards, providing strong evidence that virulent populations were already present on *Malus* species acting as reservoirs of virulence (Leroy et al., 2016). Here, we demonstrate that all identified AvrRvi6 virulent alleles originate from the wild compartment which has also been shown for some virulent isolates of *Uromyces beticola* on beet (McMullan et al., 2025).

Dating of the polymorphism at the *AvrRvi6* locus showed that five identified virulent alleles are the result of ancient non-synonymous mutations, deletion and insertion of a MITE element in the promoter. Surprisingly, no contemporary loss of avirulence event through mutation has then occurred at this locus in orchards, despite the widespread cultivation of *Rvi6* varieties exerting selection pressure on the population across Europe. This absence of new virulent alleles might be linked to the difficulty for strains to lose function likely associated to this locus in an agricultural environment, a constraint that did not appear to exist tens of thousands of years ago on wild apple trees in forest ecosystems. It is also possible that the relatively recent deployment of *Rvi6* cultivars in Europe did not exert sufficient selective pressure on local pathogen populations. The observation in transient expression experiments that AvrRvi6 alleles showed great variability in their HR-inducing capacity, is coherent with *Rvi6* not imposing a strong selection. However, the commercial success of *Rvi6*-based apple cultivars, such as Story in southern Europe, is likely to intensify selective pressure, thereby potentially creating conditions conducive to the emergence and fixation of novel virulence alleles.

Taken together, we argue that the deployment of *Rvi6* cultivar has favored the introduction of scab populations from wild *Malus* species into orchards, without necessarily promoting the emergence of new virulent alleles, as observed in many other crops (Daverdin et al., 2012; McDonald and Linde, 2002). Finally, the detection of identical *AvrRvi6* alleles in USA and Europe strongly supports the hypothesis that the virulent population was also carried along during the transatlantic spreading of apple scab since the 16th century from Europe to USA (Gladieux et al., 2008).

### AvrRvi6 roles during host colonization

The virulence function of fungal effectors is often difficult to determine as single effector deletions usually have no measurable impact on virulence as shown in *Colletotrichum orbiculare*, *P. oryzae*, and *Ustilago maydis* (Ali et al., 2014; Inoue et al., 2023; Saitoh et al., 2012). The analysis of the 122 isolates showed that 60% carry non-functional mutations which suggest that either *AvrRvi6* is not essential for the virulence of *V. inaequalis* or *AvrRvi6* function is compensated in these strains. The ectopic expression of *AvrRvi6^EU-B04^* in the virulent 2828 strain carrying the truncated allele did not bring any measurable increase in virulence on susceptible cultivars (Figure 2). Similarly, in *M. oryzae* the ectopic expression of the MAX effector *AVR1-CO39* did not contribute to enhanced virulence (Ribot et al., 2013). It is possible that *AvrRvi6* has a role during another part of the life cycle or in a specific context that was not measured in our controlled environment such as an interaction with the orchard microbiota (Snelders et al., 2022). The extensive AvrRvi6-paralogs repertoire in *V. inaequalis* (75 paralogs in the isolates MNH120 and EU-B04) raises the possibility of functional redundancy, whereby other related genes may partially compensate for the loss of *AvrRvi6* (Rocafort et al., 2022). Such redundancy has been observed in other filamentous plant pathogens, where effector families often display overlapping roles in host colonization (Raffaele and Kamoun, 2012). However, the low percentage of sequence identity between *AvrRvi6*-like effectors beyond the structural core suggest different surface properties potentially reflecting independent virulence functions (Lahfa et al., 2024). In both the MNH120 and EU-B04 isolates, *AvrRvi6* is not the most highly expressed MAX effector during subcuticular colonization, suggesting that some *AvrRvi6* paralogs play a role in the biotrophic infection of apple tissues (Supplemental Table 3). Further functional characterization of *V. inaequalis* MAX effectors is needed to fully comprehend the virulence function of this family.

### Robustness of Rvi6-mediated detection

Our allelic survey of *AvrRvi6* across a large panel of *V. inaequalis* isolates revealed substantial polymorphism, yet most variants retained the ability to trigger *Rvi6*-mediated immunity. This robustness indicates that Rvi6 recognition is highly resilient to effector diversification. Nevertheless, the strength of the immune response varied among AvrRvi6 alleles in transient expression assays, suggesting that naturally occurring polymorphisms can modulate the efficiency of *Rvi6*-mediated recognition. The biological basis of this variation remains unclear, as differences in HR intensity could reflect changes in the interaction between *Rvi6* and AvrRvi6, altered effector stability or accumulation, or differences in the efficiency of downstream immune activation. Interestingly, the avirulent alleles displayed considerable protein sequence diversity, with nine nonsynonymous substitutions compared with only a single synonymous substitution identified in isolate 3035 (Supplementary Figure 5). Although nonsynonymous variation is generally expected to be more constrained than synonymous variation under purifying selection, its presence across multiple avirulent alleles suggests that substantial sequence diversification is tolerated while preserving Rvi6 recognition. Some of these polymorphisms may nevertheless affect the efficiency of recognition without being sufficient to abolish it. Among the mutations most likely to mediate escape from recognition, we identified a small number of high-impact substitutions. These include two cysteine substitutions, C21S and C50R, which are predicted to disrupt the first and third disulfide bonds, respectively. The first disulfide bond is conserved across fungal MAX effectors identified to date and is likely important for proper protein folding (Lahfa et al., 2024). Another notable substitution, F36S, found in 15 isolates, replaces a bulky hydrophobic aromatic residue with a small polar residue and could substantially alter the local structure or surface properties of the effector. In addition, virulent alleles were sometimes associated with truncated AvrRvi6 sequences or complete loss of expression, strategies commonly observed in other gene-for-gene systems, including transposon insertion in the promoter of AvrPita1 in *P. oryzae* (Kang et al., 2001) and deletions of AvrLm1 and AvrLm6 in *Leptosphaeria maculans* (Fudal et al., 2009; Gout et al., 2007).

### AvrRvi6 is a MAX effector acting in the apoplast

AvrRvi6 belongs to the MAX effector family, a structurally defined class of fungal effectors originally described in *P. oryzae* (de Guillen et al., 2015). The MAX fold is now recognized across a wide range of Ascomycota fungi, including both Sordariomycetes and Dothideomycetes (Derbyshire and Raffaele, 2023; Le Naour-Vernet et al., 2025; Seong and Krasileva, 2023). In *V. inaequalis*, the MAX effector family has undergone a remarkable expansion, with 75 members identified in the MNH120 and EU-B04 genomes, making it the largest effector family in this pathogen. To date, all the characterized MAX effectors are translocated into the host cytoplasm to manipulate intracellular immune signaling (Khang et al., 2010; Ribot et al., 2013; Sornkom et al., 2017). Our work reveals that MAX effectors can also act extracellularly in the apoplast where they could fulfill similar functions in manipulating the plant immune system. Structural analyses revealed that *V. inaequalis* MAX proteins, including AvrRvi6, contain additional disulfide bonds compared to their *P. oryzae* counterparts (Lahfa et al., 2024). We propose that these extra bonds enhance protein stability, enabling survival in the harsh, protease-rich apoplastic environment. This hypothesis aligns with studies of other apoplastic effectors, where structural stabilization is a common feature conferring resistance to host proteases (Rocafort et al., 2020; Sperschneider et al., 2018). In *P. oryzae*, the MAX family is enriched for avirulence factors, suggesting that plants have repeatedly evolved immune receptors to detect this conserved fold (Le Naour-Vernet et al., 2025). The fact that the first cloned *AVR* gene in *V. inaequalis* is a MAX effector suggests that other members of this family may also act as recognition targets for apple immune receptors. Given the large size of the MAX repertoire in *V. inaequalis*, and that many of the genes encoding these proteins are highly expressed during host colonization, future studies are likely to uncover additional *R* gene-effector pairs, providing a roadmap for durable resistance breeding.

### Concluding remarks and future perspectives

By identifying and characterizing *AvrRvi6*, we provide the first molecular insight into the interaction between *V. inaequalis* and its apple host. The unexpected discovery that a MAX effector functions in the apoplast challenges current paradigms and suggests that this structural family is even more versatile than previously appreciated. Looking ahead, comprehensive functional characterization of the *V. inaequalis* MAX repertoire will be essential for fully understanding the molecular basis of this interaction and for informing future breeding strategies. From a practical perspective, our study provides two key contributions to resistance management. First, we show that virulence against *Rvi6* has been present for thousands of years in wild *Malus* species across the northern hemisphere, with no evidence of *AvrRvi6* alleles that emerged following the deployment of resistant cultivars. Because no *Malus* species are naturally endemic to the southern hemisphere, we argue that *Rvi6*-based resistance could remain durable there, provided that strict measures are taken to prevent the introduction of virulent strains from the northern hemisphere. Second, as no recent mutational events leading to virulence have been detected, a molecular tool targeting all virulent polymorphisms in the *AvrRvi6* locus and its promoter could be used to monitor virulent populations, thereby guiding the deployment of resistant cultivars and reducing the risk of resistance breakdown.

## Materials and Methods

### Fungal strains and culture conditions

Three sets of *Venturia inaequalis* isolates were used in the present study: (i) 51 progeny from an *in vitro* cross between the avirulent isolate 0301 and the virulent isolate 1066 ((Bénaouf and Parisi, 2000); Supplemental Table 1)), (ii) 41 isolates sampled in an orchard with recombination between avirulent and virulent populations towards *Rvi6* (Supplemental Table 1), and (iii) 122 isolates sampled in 21 different countries worldwide (Supplemental Table 5), most of them from *Malus × domestica* on cultivars with or without *Rvi6*. Other isolates came from different *Malus* species, especially *M. floribunda* (source of *Rvi6* resistance), *M. sieversii* and *M. sylvestris* (progenitors of *M. × domestica*). All isolates were cultured on cellophane membranes (400P, Jacques Emballage) placed on PDA-YE (Potato Dextrose Agar 39 g L^-1^ with Yeast Extract 3 g L^-1^) medium at 18–20 °C under a 16 h light/8 h dark photoperiod, as described by (Caffier et al., 2014). The use of cellophane membranes facilitates improved production and harvesting of conidia.

### Apple pathogenicity assays

Pathogenicity tests were conducted using apple trees of *M. × domestica* grafted onto the rootstock MM106. Apple cultivars Gala (X04712) and Golden Delicious (X00972) were used as controls for susceptibility, whereas apple cultivars J45 (Caffier et al., 2014) and Priscilla (H6, (Bus et al., 2011)), which carry the *Rvi6* resistance gene against *V. inaequalis*, were used to evaluate the virulence of the isolates against this specific resistance. Plants were grown in a glasshouse in a scab-free environment and transferred into a growth chamber for pathogenicity tests. For each *V. inaequalis* isolate, spores were harvested by shaking the cellophane membranes in sterile water. After filtration through medical gauze, the conidial suspensions were adjusted to a final concentration of 2.5 × 10⁵ spores / mL. *V. inaequalis* isolates were inoculated onto at least three plants of each apple cultivar using a manual sprayer. Following inoculation, plants were kept in darkness under a plastic sheet to maintain leaf wetness. The climatic conditions in the growth chamber were set to 95% relative humidity and 17°C for the first 48 h. After this period, the plastic sheet was removed, and the relative humidity was adjusted to 80% during the day and 90% at night, with a 12-h light period each day. To assess the ability of each isolate to cause disease, the presence or absence of scab symptoms on each plant was recorded at 14 and 21 days post-inoculation.

### Genomic DNA extraction for sequencing

*V. inaequalis* isolates were grown on cellophane membranes as described above. Mycelium and spores were scraped from the surface of cellophane membranes and fast-frozen in liquid nitrogen prior to grinding. Genomic DNA was extracted from 100 mg of ground mycelium using the Nucleospin Food kit (740945.50, Macherey Nagel), according to the manufacturer’s recommendations. Quality and quantity of the DNA was measured using a NanoDrop One (Thermo Scientific).

### Library preparation, pool sequencing, read mapping and SNP calling

Genomic DNA from *V. inaequalis* isolates was pooled according to both origin and virulence phenotype, generating four pools: progeny-avirulent, progeny-virulent, orchard-avirulent, and orchard-virulent isolates. These four pools were used to generate paired-end genomic DNA libraries. Libraries were then sequenced on a HiSeq2500 sequencer (Illumina, San Diego, CA, USA) with 2 ×100 paired-end reads. Raw reads were trimmed using Trimmomatic v. 0.32 (Bolger et al. 2014) with a sliding-window approach, in which reads were trimmed when the mean quality score within a 4-bp window fell below 15. Low quality bases (<20) from both ends and reads shorter than 36 nucleotides were also removed. The remaining reads were mapped onto the *V. inaequalis* reference genome assembly EU-B04 (Le Cam et al., 2019) with bowtie2 v. 2.1.0 (Langmead et al., 2009) using standard parameters for the “sensitive end-to-end” mode. PCR duplicates were removed using Picard v. 1.88 (http://broadinstitute.github.io/picard/). Biallelic SNPs were called using Samtools v.1.1 (Li, 2011) and Popoolation2 v. 1.201 (Kofler et al., 2011). Although the SNP calling was performed separately for genetic variation found in natural populations and SNPs segregating in the progeny, a similar strategy was employed. First, SNPs with at least 10 reads on the alternate alleles, a depth per pool at the population greater than 50X but lower than 300X were identified. Second, to minimize false positive calls due to Illumina sequencing errors, all SNPs with a minor allele frequency (MAF) lower than 0.05 in natural populations and 0.1 in the progeny were discarded using a custom script available upon request (available for download at a GitHub repository). A total of 5,254,356 and 3,104,118 SNP loci were identified in the natural populations and progeny, respectively. Sliding window Fst estimates were then computed from allele frequencies derived from allele counts using the standard method implemented in fst-sliding.pl, available in the popoolation2 bioinformatics suite (Kofler et al., 2011).

### RNA-seq analysis

Detached leaf infection assays were carried out with the isolate EU-B04 on the youngest leaves of apple cultivar Gala. Single leaves were placed on Petri dishes containing water-saturated tissue paper and the cut end of the petioles were covered with damp tissue paper (adaxial side up). The leaves were inoculated with 5 µL droplets of conidia suspension (1.5 x 10^5^ spores per mL) and incubated for 48 h in the absence of light. Three biological replicates were prepared for each isolate consisting of three infected leaves each. Infected leaf material was collected 48 h after inoculation and frozen in liquid nitrogen and ground to a fine powder using a bead beater. Total RNA was extracted from the ground material following the protocol described by (Chang et al., 1993) and the samples were treated with DNAse. Sequencing libraries were prepared with the TruSeq LT kit and RNA sequencing was performed on a High-Seq 3000 (2 × 150 paired end reads). Sequencing reads were first quality trimmed using Trimmomatic (v. 0.39, (Bolger et al., 2014)) with the recommended parameters for paired-end reads. The processed reads were then mapped to the *V. inaequalis* EU-B04 reference genome (Le Cam et al., 2019) using the splice-aware aligner HISAT2 (v. 2.2., (Kim et al., 2019)). Resulting sam files were converted to bam format and sorted using SAMtools (v. 1.21, (Li et al., 2009)). Gene expression quantification was based on the gene models established for isolate EU-B04 (Le Cam et al., 2019). The gene model file was first converted to a GFF3 file in which we reconstructed the hierarchy between gene mRNA and exon/CDS, using AGAT (v. 1.2.0 (Dainat, 2022)) and the script *agat_convert_sp_gxf2gxf.pl*. The resulting GFF3 file was then converted to GTF format using gffread (version 0.12.6, (Pertea and Pertea, 2020)). Raw gene expression counts were computed using HTSeq-count (v. 2.0.4, (Anders et al., 2015)) using the EU-B04 gene model as reference. Raw counts were finally normalized using Transcripts Per Million (TPM) method to allow proper comparison between replicates. To do so, gene lengths were first computed using the tool “Gene length and GC content from GTF and FASTA file” from the European instance of Galaxy (Galaxy v. 0.1.2, (Giardine et al., 2005; Goecks et al., 2010)). Using this gene length information, raw counts were converted to TPM using the “convertCounts” function from “DGEobj.utils” R package (Law et al., 2018) under R v. 4.5.1 (R Core Team 2025). Unless stated otherwise, analyses were performed on the Genotoul bioinformatics platform Toulouse Occitanie (Bioinfo Genotoul, https://doi.org/10.15454/1.5572369328961167E12).

### Plasmid construction

For the transformation of *V. inaequalis*, the binary vector pJK5, suitable for *Agrobacterium tumefaciens*-mediated transformation was modified by inserting the *AvrRvi6* gene, under the control of its native promoter. The selectable marker was the *hygromycin B phosphotransferase* gene (*hph*), driven by the *Aspergillus nidulans OliC* promoter. For the transient transformation of *N. benthamiana,* constructs were generated using the Golden Gate Modular Cloning (MoClo) Toolkit, plant part and plant part II kits (Engler et al., 2014; Gantner et al., 2018; Weber et al., 2011). The tested alleles *AvrRvi6^ΔSP^* or *Avr4 ^ΔSP^* were cloned in a L0 CDS1 non-stop module and assembled with the *2×35S promoter*, *SP^PR1α^*, *mCherry* or *6×HA* and 35S terminator in the L1 binary vector pICH47811. Similarly, full length *Rvi6* or *Cf-4* coding sequences were cloned in L0 CDS1 non-stop module and assembled with the 2×35S promoter and 35S terminator in the L1 binary vector pICH47811. Cloning design and sequence analysis were done using Geneious Prime v9.0.5. All constructs were confirmed by restriction enzyme digestion and Sanger sequencing prior to use in transformation. Plasmid construction is described in Supplemental Table 6.

### Agrobacterium tumefaciens-Mediated Transformation of V. inaequalis

Strain 2828 of *V. inaequalis*, known for its virulence and ability to overcome *Rvi6* resistance in host plants, was used. After 14 days of incubation, the fungal material was collected by carefully scraping the surface of the cellophane membrane with a sterile scalpel. This material was then transferred into sterile water and homogenized using a hand-held blender. The grinding process facilitates the efficient release of conidia and other fungal structures, enhancing the overall transformation efficiency. Transformation was performed using *Agrobacterium tumefaciens* strain LBA4404 containing the binary vector, which was introduced into the bacterial strain by electroporation. *A. tumefaciens* was cultured overnight in Yeast Extract Mannitol (YM) medium with rifampicin (50 µg/mL), streptomycin (50 µg/mL) and chloramphenicol (25 µg/ml), then diluted to an OD₆₀₀ of 0.15 in induction medium (IM) supplemented with 200 µM acetosyringone. The bacterial culture was incubated at 28 °C for 5 h with shaking. Conidial suspensions from *V. inaequalis* strain 2828 were mixed with the induced *A. tumefaciens* culture in a 1:1 ratio. The mixture was placed onto sterile cellophane membranes on IM agar plates and co-cultivated for 60 h at 24 °C in the dark. After co-cultivation, membranes were transferred to PDAYE plates supplemented with hygromycin B (30 µg/mL) and cefotaxime (200 µg/mL) to eliminate residual bacteria. Plates were incubated at 20 °C for up to 4 weeks, and hygromycin-resistant colonies were isolated and subcultured onto fresh selective media. Confirmation of transformed strains was performed by extracting genomic DNA from hygromycin-resistant colonies using the NucleoSpin Food kit (Macherey-Nagel). Transgene integration was confirmed by PCR amplification of both the *hph* gene and the gene of interest. The phenotypic characteristics of the transformants were compared with those of the wild-type isolate 2828 to evaluate morphological stability *in vitro* and pathogenicity on susceptible and *Rvi6* cultivars. Pathogenicity assays were carried out in a confined growth chamber, using the same protocol as described above. The pathogenicity of each isolate on each apple plant was evaluated quantitatively through visual estimation of the percentage of diseased leaf area on five leaves per plant. Mean disease area were analyzed with a one-way ANOVA at a significance level of *P* < 0.05 followed by a multiple comparisons of means using Tukey’s post hoc test.

### Agrobacterium tumefaciens-mediated transformation of N. benthamiana

*A. tumefaciens* cultures were grown for 48 h at 28°C on LB plates with the corresponding antibiotics (Supplemental table 6.). After which, the cultures were resuspended in 2 mL of induction medium (50 mM MES, pH 5.6, 0.5% [w/v] and 200 mM acetosyringone) and incubated in the dark for 4 h. Each culture’s optical density at 600 nm (OD_600_) was then adjusted to 2.0. The cultures containing each plasmid were mixed in equal volumes to achieve a final OD_600_ of 0.25 per construct. The abaxial leaf surface of 5- to 7-week-old *N. benthamiana* plants were hand-infiltrated with a needleless syringe. Plants remained in the same glassehouse condition with ∼20 °C and 16 h photoperiod for 4 additional days before phenotyping.

### Hypersensitive response (HR) assay

Following the transformation of *N. benthamiana* to express constructs of interest (Supplemental Table 6), quantification of HR intensity was performed using red-light imaging on a Chemidoc apparatus (Biorad) as described in (Landeo Villanueva et al., 2021). Median pixel intensity was measured using FIJI (Schindelin et al., 2012) and then normalized against the HR produced by the co-expression of *Cf-4* and *Avr4* present on each leaf. Three *AvrRvi6* alleles plus *Avr4* were tested in co-construction with *Rvi6* and *Cf-4* on each leaf (three biological replicates for each allele on three different *N. benthamiana* leaves). The experiment was done twice. The effect of alleles on the normalized pixel intensity was analyzed using an ANOVA with two factors (*AvrRvi6* allele and Experiment) for samples co-transformed with *Cf-4* and *Rvi6* separately. The relative pixel intensity was submitted to Log transformation in order to respect the conditions of normality and homoscedasticity of the residuals. When a significant effect of alleles was observed, a comparison of each allele with the reference *Avr4* was performed using a Dunnet test.

### Laser-scanning confocal microscopy

*N. benthamiana* leaves transiently expressing the constructs of interest were sampled 3 days post-agroinfiltration. Leaf fragments were infiltrated with water then mounted on microscope slides. Imaging was performed using Nikon AX laser-scanning confocal microscope (IMAC platfrom, SFR QuaSav, Angers) equipped with a 40X oil-immersion objective (Plan Fluor 40X oil DIC H N2). The fluorescence of mEGFP, mCherry and chlorophyll was observed using the following excitation/emission wavelengths: 488/499-533, 561/592-625 and 640/662-737 respectively. Images were processed using Fiji software (Schindelin et al., 2012). The fluorescence intensity plots were generated using the plot profile tool in Fiji to quantify pixel intensity along a line for each channel (GFPand mCherry). The dataset was normalized based on the maximal fluorescence intensity values.

### Western blot analysis

Western blots were performed on either apoplastic wash fluid or crude protein extract. For apoplastic wash fluid extraction, transformed *N. benthamiana* leaves were vacuum infiltrated with ice-cold water. Excess water was removed using absorbent paper, and leaves were placed in an empty 20 mL syringe inserted into a 50 mL Falcon tube. Apoplastic wash fluid was collected by centrifugation at 2,000 × g for 10 min at 4°C and immediately flash-frozen in liquid nitrogen. For crude protein extraction, a 1.5 cm² leaf disc from agroinfiltrated *N. benthamiana* leaves was ground directly in liquid nitrogen. Apoplastic wash fluid samples and ground leaf tissue samples were supplemented with Laemmli buffer and heated at 95°C for 5 min. Proteins were separated on 12% SDS-polyacrylamide gels and transferred to membranes for immunoblot analysis using an anti-hemagglutinin (HA) antibody (Roche, 12013819001) or an anti-red fluorescent protein (RFP) antibody (Rockland, 600-401-379).

### Evolutionary analyses

To explain the polymorphism observed in the *AvrRvi6* gene region, we used the SNP information from 17 individuals of *V. inaequalis* belonging to the *Rvi6*-virulent population and 21 individuals from the *Rvi6*-avirulent population, both sampled on European domestic apple trees (Supplemental Table 5). For this, we used the coalescent-based method implemented in Relate (Speidel et al., 2019). Population demographic dynamics and split time were first estimated in order to test for a split subsequent to the *Rvi6* gene deployment, or a more ancient divergence time as previously suggested by (Lemaire et al., 2016). As virulent alleles might have been selected in resistant apple trees, selection at the region encompassing the *AvrRvi6* gene was also tested. This selection test is based on the speed of increase in frequency of a particular lineage carrying a given mutation compared to all other lineages (Speidel et al., 2019). Lastly, the genealogical framework proposed by Relate allows us to estimate the age of mutations, and especially those responsible of the virulent phenotype. For all Relate analyses, a mutation rate of 3.3×10^-10^ (the haploid mutation rate of yeast *Saccharomyces cerevisiae* (Lynch et al., 2008), a generation time of 1 (one obligatory sexual event per year) and a total of 10 iterations were chosen. For each analysis, three repetitions were performed to ensure convergence.

### Identification and characterization of *AvrRvi6*-like sequences

To identify *AvrRvi6*-like sequences in *V. inaequalis* isolate EU-B04, a tBLASTn search was performed against the EU-B04 genome using the AvrRvi6 protein and any retrieved homologs as queries in Geneious v9.0.5 (Kearse et al., 2012). Where possible, exon–intron boundaries of the resulting *AvrRvi6*-like sequences were inferred using RNA-seq data from EU-B04 (this study) or from isolate MNH120 (Rocafort et al., 2022). Sequences were considered full-length if they met all of the following criteria: (1) the N- and C-termini were intact, (2) no premature stop codons or frameshift mutations were present, (3) canonical GT–AG intron donor–acceptor splice sites were present, and (4) all six conserved cysteine codons were present. Signal peptides were predicted in AvrRvi6-like proteins using SignalP v4.1 (Nielsen, 2017) and aligned using Geneious v 9.0.5.

## Acknowledgements

We thank Isabel Guerrero-Zepeda, Louis Amossé, Ophélie Dubreu and Nicolas Anger for assisting with some of the experimental procedure. The Biological Resource Center “RosePom - Pome Fruits and Roses” (https://irhs.angers-nantes.hub.inrae.fr/dispositifs-mutualises/ressources-genetiques/crb-fruits-a-pepins-et-rosier) and the Horticulture Experimental Facility UE HORTI (https://doi.org/10.15454/1.5573931618268674E12) are acknowledged for maintaining apple material in orchard and the Angers Plant Phenotyping facility PHENOTIC (https://doi.org/10.17180/ykbz-2v85) is acknowledged for the production of plants in glasshouses. We thank the IMAC (Cellular Imaging) technical platform of SFR 4207 QUASAV, Université d’Angers. We thank Stella Césari for pointing out that AvrRvi6 looks like a MAX effector and Benjamin Petre for the kind gift of the GFP-PITG16242 plasmid. We thank the many contributors who sent isolates of *V. inaequalis* from around the world.

## Author contributions

Conceptualizations: JC, CL, BLC, MB

Methodology: MS, JC

Investigation: MS, SB, JC, CL, VC, CD, JS, EMB, AV, HS, CM, BLC, MB

Funding acquisition: CL, BLC, MB

Writing - original draft: MS, SB, JC, CL, VC, CM, BLC, MB

Writing – review & editing: SB, JC, VC, EMB, CM, BLC, MB

## Data and materials availability

Whole genome sequencing and RNAseq data have been deposited on the NCBI BioProject PRJNA1391408 (https://www.ncbi.nlm.nih.gov/bioproject/1391408) and will be released upon publication. All plasmids, and fungal strains generated in this work are available from the authors upon request. Phenotypic data on virulence of *Venturia inaequalis* strains have been deposited on the Recherche Data Gouv Entrepot (https://doi.org/10.57745/BZR9DU).

## Supplemental table and figure legends

Supplemental Table 1: Description of pooled genome sequencing data.

Supplemental Table 2: RNAseq of isolate EU-B04 inoculated on Gala detached leaves 48 hours post-inoculation. Normalized expression levels (TPM) are shown for three independent biological replicates.

Supplemental Table 3: *AvrRvi6*-like sequences in the genome of *Venturia inaequalis* isolate EU-B04. Introns are shown as red text. In some cases, intron sequences could not be predicted. For each nucleotide sequence, the corresponding genome scaffold position and predicted EU-B04 gene number (where available) are provided. Predicted protein sequences are shown next the nucleotide sequences (in some cases, mutations prevented prediction). Signal peptides are underlined. The average expression (Transcripts Per Million) and standard deviation from 3 independent replicates is shown (where available).

Supplemental Table 4: Percentage amino acid sequence identity between AvrRvi6 and AvrRvi6-like sequences from *Venturia inaequalis* isolate EU-B04.

Supplemental Table 5: List of *Venturia inaequalis* isolates used in this study.

Supplemental Table 6: Description of plasmid constructs.

Supplemental Figure 1: Molecular validation of *Venturia inaequalis* transformants used in Figure 2. Agarose gel electrophoresis of PCR products with specific primers on the T-DNA constructions in *V. inaequalis* isolate 2828. Lane 1: 1 kb Plus DNA Ladder (NEB); lane 2: wild-type (WT) isolate 2828; lane 3: 2828 isolate transformed with GFP construct; lanes 4–5: two independent 2828 transformants expressing *AvrRvi6^EU-B04^* (AvrRvi6 #1 and #2); lane 6, plasmid pJK5 (positive control); lane 7: water control (H₂O). Molecular sizes (kb) are indicated on the left.

Supplemental Figure 2: Expression of *AvrRvi6^EUB-04^* induces *Rvi6*-dependant resistance. Representative phenotypes are shown for each isolate on susceptible cultivars Gala and Golden and on *Rvi6*-carrying cultivars J45 and Priscilla. The number of plants with disease over the number of plants inoculated is shown below each leaf. Bar = 1cm

Supplemental Figure 3: Protein alignment of AvrRvi6-like proteins. Identity percentage for each amino acid position is indicated on top.

Supplemental Figure 4: AvrRvi6 sequence diversity in *Venturia inaequalis* strains. Nucleotide and protein alignment of AvrRvi6 coding sequences from 122 strains of *Venturia inaequalis*. Polymorphisms associated with virulence are framed in red.

Supplemental Figure 5: Virulent isolates carrying the *AvrRvi6^0301^* allele have a different promoter associated with reduced expression during infection. **A.** Nucleotide alignment of the *AvrRvi6* promoter in different isolates carrying the *AvrRvi6^0301^* allele. Start codon is highlighted in green, 100% consensus in red and 50% consensus in blue. **B.** Annotation of a miniature inverted-repeat transposable element (MITE) in the promoter of *AvrRvi6* in isolate 1066. Terminal inverted repeats (TIR) are highlighted in pink **C.** Relative expression of *AvrRvi6* in isolates 3040 and 1066 at 3, 7 and 15 days post inoculation.

Supplemental Figure 6: Selection analyses of *AvrRvi6* alleles. **A-B.** Selection analysis in the *AvrRvi6* region of the virulent population on a portion of chromosome 18 (A) or the entire chromosome 18 (B). **C.** Ages of branches carrying the mutations on chromosome 18. **A-C.** The *AvrRvi6* region (1kb around) is indicated in red.

