## Supplemental Figure for "Ancient MAX Effector Variants of a Fungal Pathogen Evade Apoplastic Immunity in Apple"

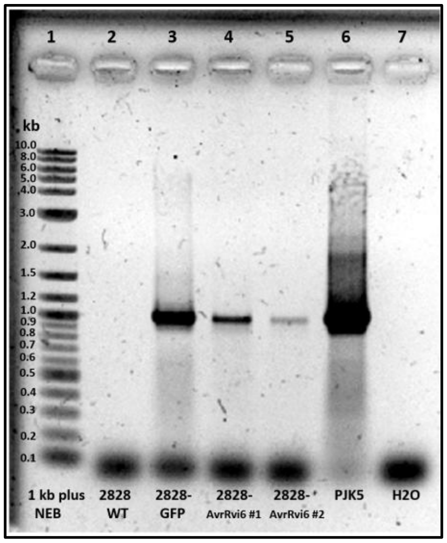


Supplemental Figure 1: Molecular validation of *Venturia inaequalis* transformants.

Agarose gel electrophoresis of PCR products with specific primers on the hygromycin gene (expected size = 1 kb) of the T-DNA constructions in *V. inaequalis* isolate 2828. Lane 1: 1 kb Plus DNA Ladder (NEB); lane 2: wild-type (WT) isolate 2828; lane 3: 2828 isolate transformed with GFP construct; lanes 4–5: two independent 2828 transformants expressing *AvrRvi6*$^{EU\text{-}B04}$ (AvrRvi6 #1 and #2); lane 6, plasmid pJK5 (positive control); lane 7: water control ($H_2O$). Molecular sizes (kb) are indicated on the left.

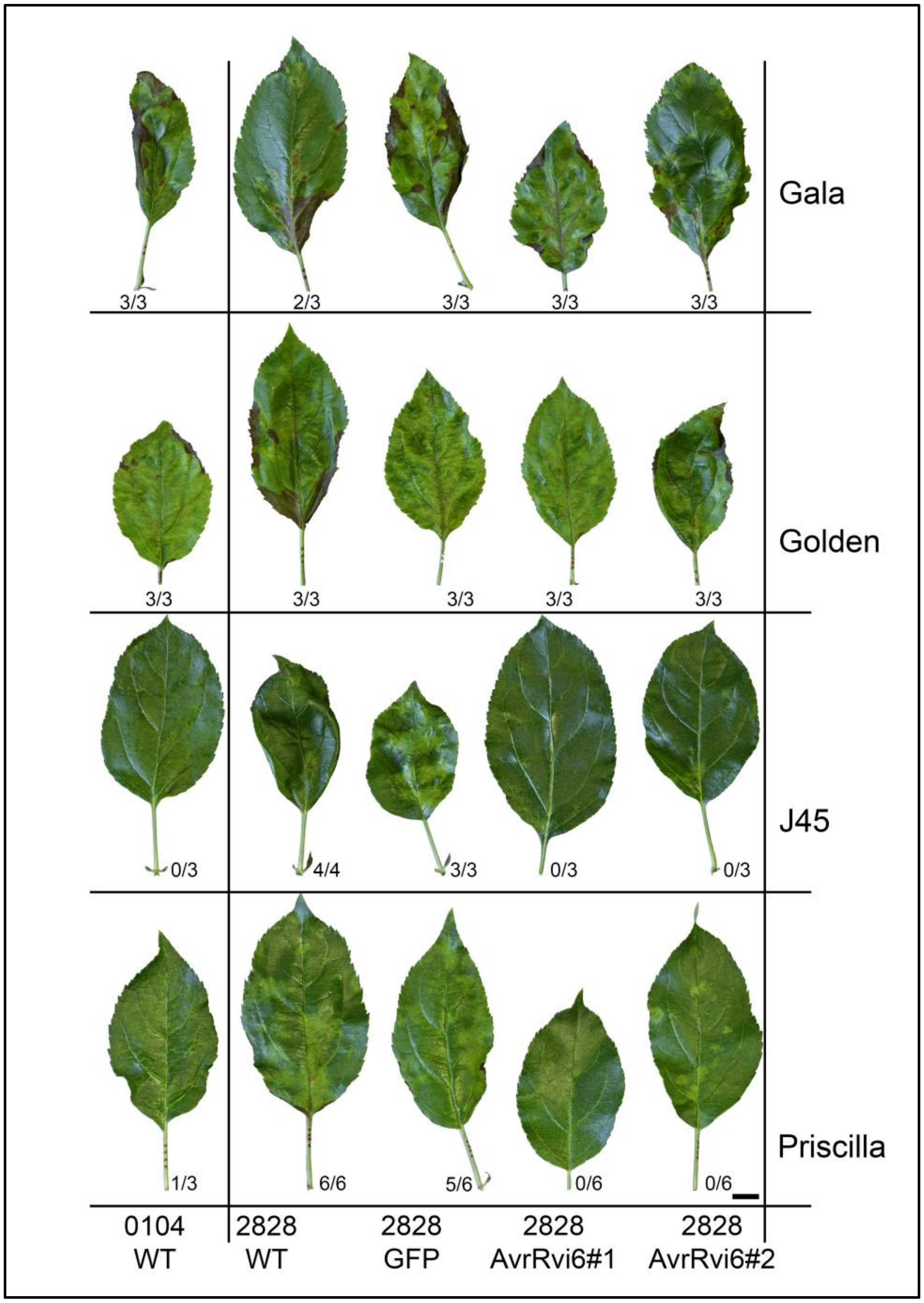


Supplemental Figure 2: Expression of *AvrRvi6*$^{EUB\text{-}04}$ induces *Rvi6*-dependant resistance. Representative phenotypes are shown for each isolate on susceptible cultivars Gala and Golden and on *Rvi6*-carrying cultivars J45 and Priscilla. The number of plants with disease over the number of plants inoculated is shown below each leaf. Bar = 1cm

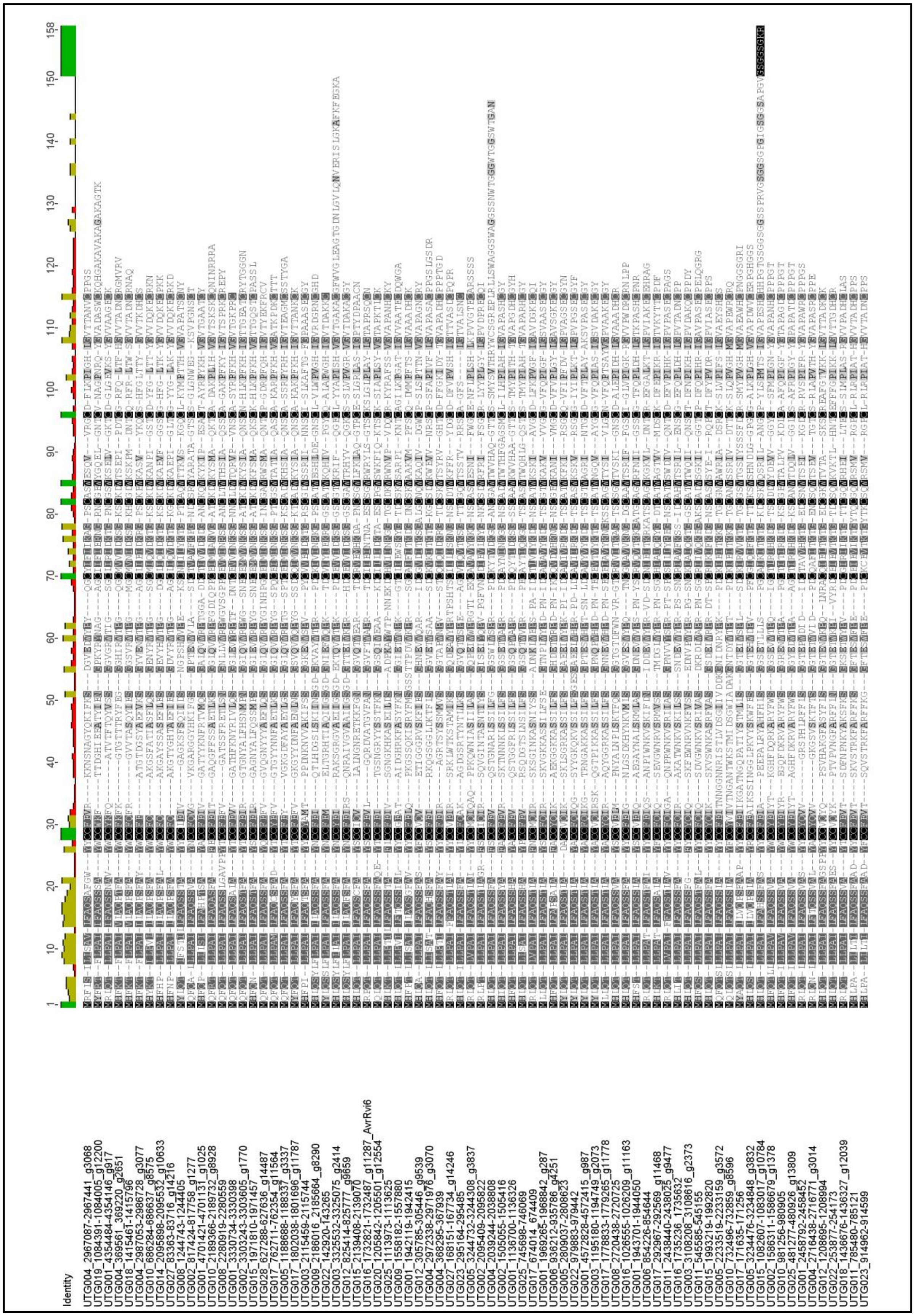

Supplemental Figure 3: Protein alignment of AvrRvi6-like proteins. Identity percentage for each amino acid position is indicated on top.

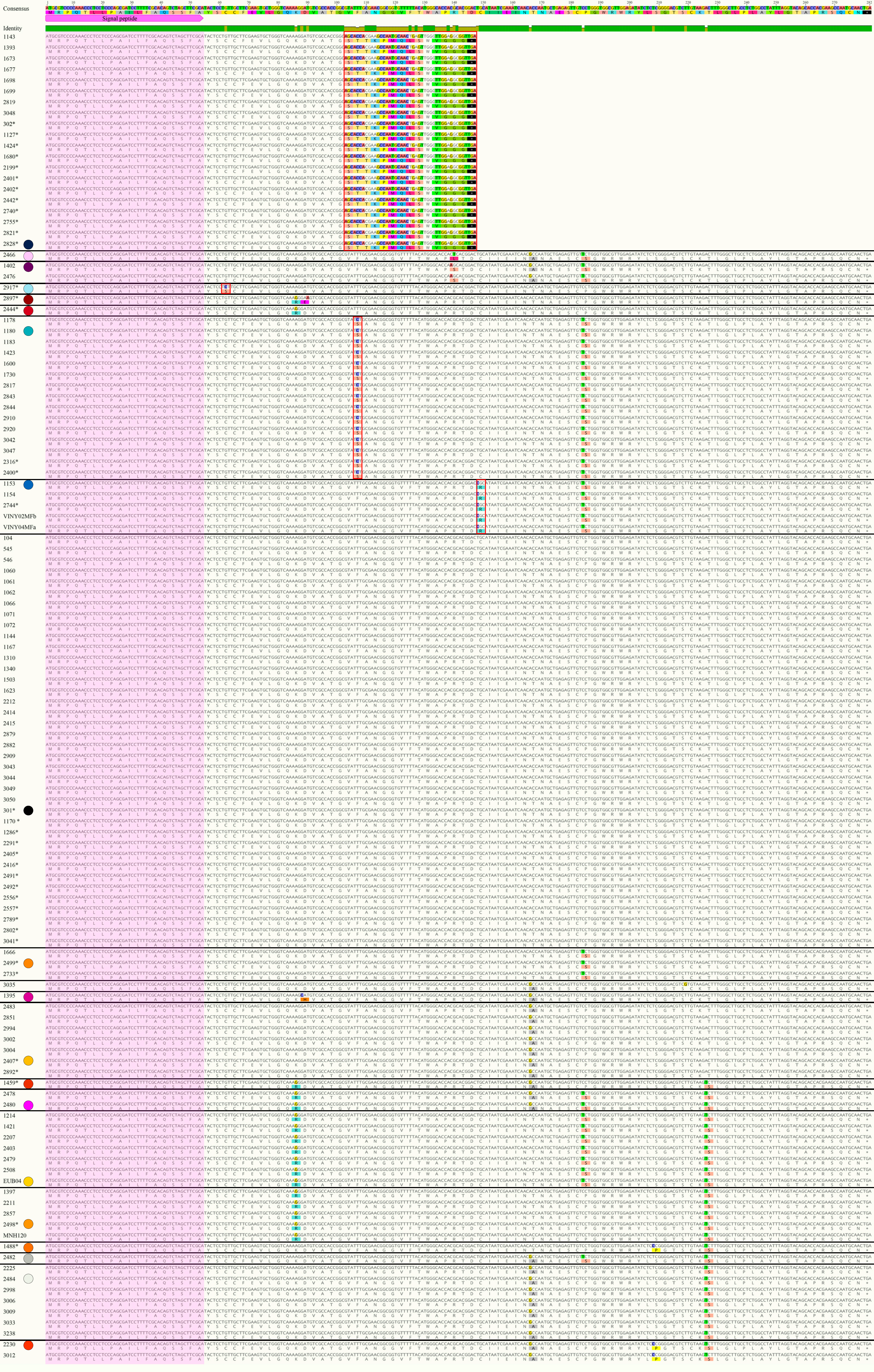

Supplemental Figure 4: AvrRvi6 sequence diversity in *Venturia inaequalis* strains.
Nucleotide and protein alignment of AvrRvi6 coding sequences from 122 strains of Venturia inaequalis. Polymorphisms associated with virulence are framed in red.

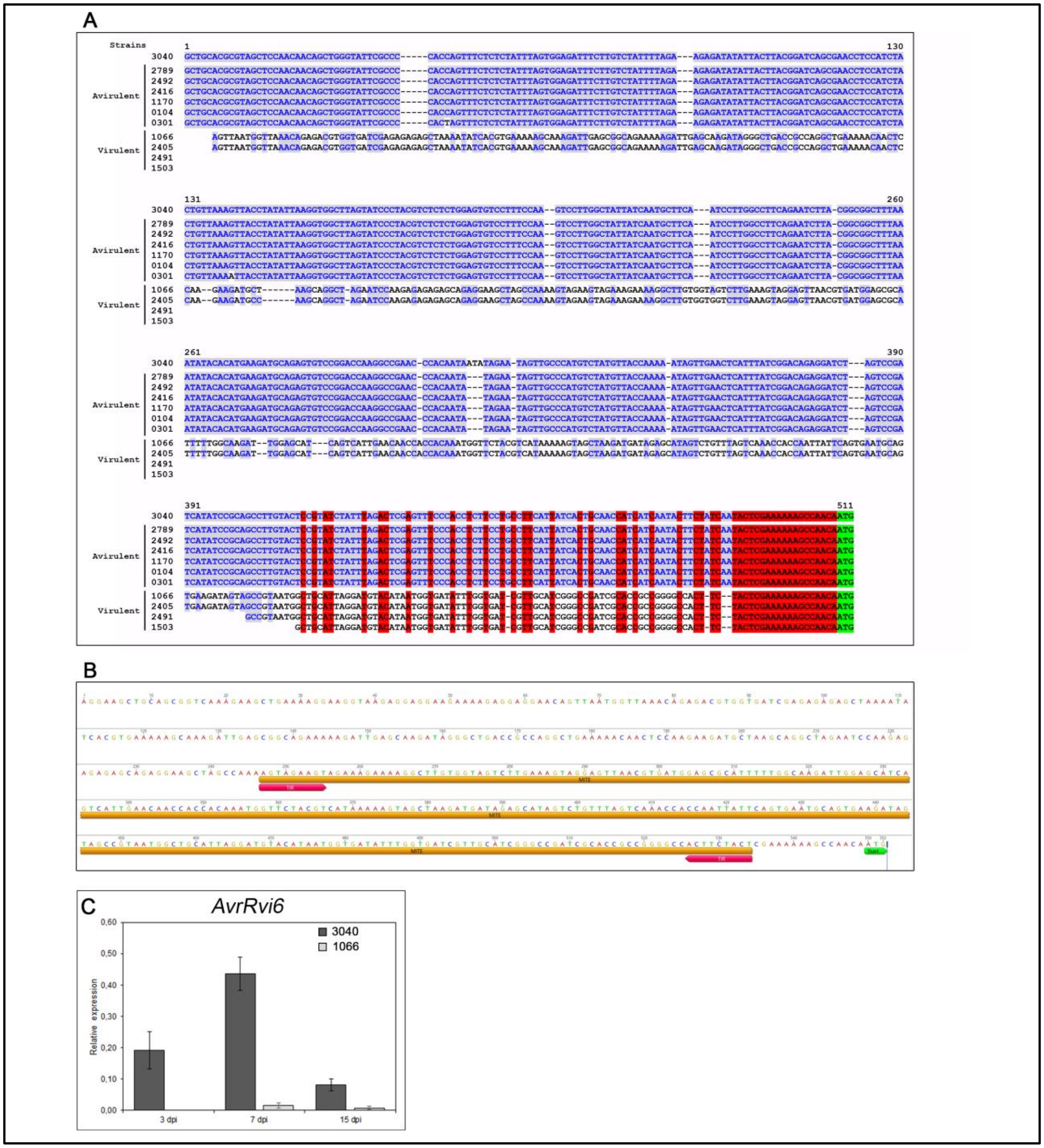


<u>Supplemental Figure 5: Virulent isolates carrying the *AvrRvi6*$^{0301}$ allele have a different promoter associated with reduced expression during infection.</u>

**A.** Nucleotide alignment of the *AvrRvi6* promoter in different isolates carrying the *AvrRvi6*$^{0301}$ allele. Start codon is highlighted in green, 100% consensus in red and 50% consensus in blue. **B.** Annotation of a miniature inverted-repeat transposable element (MITE) in the promoter of *AvrRvi6* in isolate 1066. Terminal inverted repeats (TIR) are highlighted in pink **C.** Relative expression of *AvrRvi6* in isolates 3040 and 1066 at 3, 7 and 15 days post inoculation.

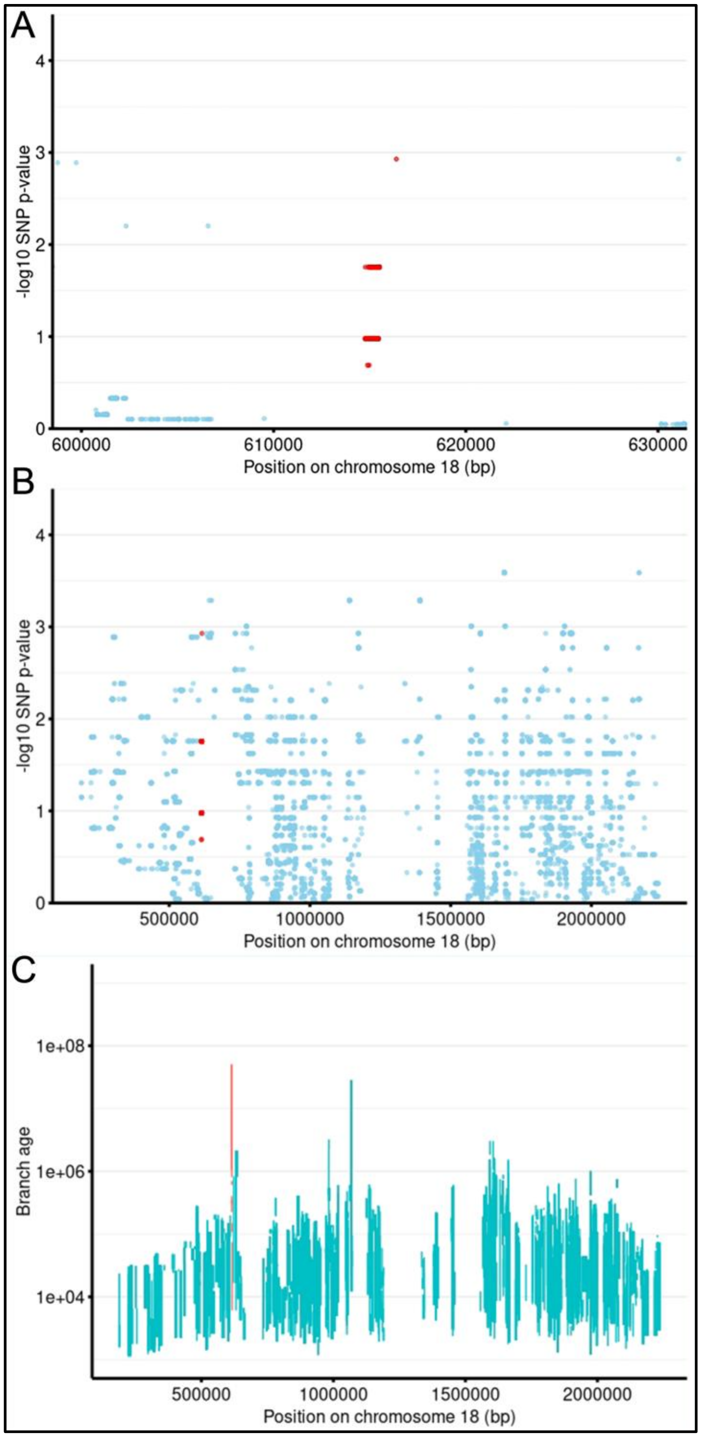


Supplemental Figure 6: Selection analyses of *AvrRvi6* alleles. **A-B.** Selection analysis in the *AvrRvi6* region of the virulent population on a portion of chromosome 18 (A) or the entire chromosome 18 (B). **C.** Ages of branches carrying the mutations on chromosome 18. **A-C.** The *AvrRvi6* region (1kb around) is indicated in red.
